# Different mechanisms link gain and loss of kinesin functions to axonal degeneration

**DOI:** 10.1101/2024.12.31.630930

**Authors:** Yu-Ting Liew, David M.D. Bailey, William Cairns, Maureece Day, Federico Dajas-Bailador, Ella Jones, Sophie McCann, Matthias Landgraf, Lydia Lorenzo-Cisneros, Thomas Murphy, Milli Owens, Devesh Pant, Jill Parkin, Haydn Tortoishell, André Voelzmann, Andreas Prokop

## Abstract

Axons are the slender, often meter-long projections of neurons that form the biological cables wiring our bodies. Most of these delicate structures must survive for an organism’s lifetime, meaning up to a century in humans. Long-term maintenance and sustained functionality of axons requires motor protein-driven transport distributing life-sustaining materials and organelles to places of need. It seems therefore plausible that loss of motor function would cause axon degeneration; however, also gain-of-function conditions were linked to disorders including motor neuron disease or spastic paraplegia. To understand this phenomenon, we studied ∼40 genetic manipulations of motor proteins, cargo linkers and regulators of reactive oxygen species in one standardised *Drosophila* primary neuron system. Using axonal microtubule bundle organisation as a relevant readout reflecting the state of axon integrity, we found that losses of Dynein heavy chain, KIF1A/Unc-104 and KIF5/Kinesin heavy chain (Khc) all cause bundle disintegration in the form of chaotically curled microtubules. Detailed functional studies of Khc and its adaptor proteins revealed that losses of mitochondrial or lysosomal transport cause ROS dyshomeostasis, which is a microtubule-curl-inducing condition in fly and mouse neurons alike. We find that hyper-activated Khc induces the same microtubule curling phenotype, not through ROS but likely more directly through enhanced mechanical forces. Studies with loss of Unc-104 or KIFBP and expression of an ALS-linked mutant form of the human Khc orthologue KIF5A suggest that loss or hyperactivation of different types of transport motors cause MT curling as a shared feature. We discuss a model which can explain our findings and their relevance for understanding motor-linked neurodegeneration.

## Introduction

Axons are the long and slender processes of neurons which form the biological cables that wire the nervous system and are indispensable for its function. In humans, axons can be up to 2 metres long at diameters of only 0.1-15 µm (Prokop, 2020). Most of these delicate cellular processes must survive for an organism’s lifetime, meaning up to a century in humans. Unsurprisingly, mammals lose about 40% of their axon mass towards high age (Calkins, 2013; Coleman, 2005; Marner et al., 2003). This rate is drastically increased in hereditary forms of axonopathies (Prokop, 2021; Smith et al., 2023).

A prominent group of axonopathy-inducing mutations affect proteins involved in live-sustaining axonal transport of RNAs, proteins, lipids and organelles, especially dynein/Dynactin, kinesin-1 and kinesin-3 family members and some of their associated adaptor proteins (Guedes-Dias and Holzbaur, 2019; Hirokawa et al., 2010; Smith et al., 2023). The loss of such transport was shown to affect axon growth, function and regeneration (Guerra San Juan et al., 2024) and can be expected to lead to systemic collapse of axonal structure and physiology, hence axonopathy. This would explain the reported links of motor protein mutations to Charcot Marie Tooth diseases, hereditary sensory neuropathies or ataxias (for a detailed list of links in the Online Mendelian Inheritance in Man^®^ database, see Smith et al., 2023). Surprisingly, disease-relevant mutations of motors do not only cause loss-of-function, but some mutations cause over-activation of motors (Baron et al., 2022; Chiba et al., 2019; Gabrych et al., 2019; Pant et al., 2022; Pino et al., 2023). This poses the question of how opposing mutational effects can cause comparable outcomes.

To address this question, we used an experimental approach based on two strategic decisions: Firstly, we used axonal microtubules (MTs) as our main readout. These MTs are arranged into loose bundles that run all along axons; these MTs are required for axonal morphogenesis and they form the essential highways for axonal transport (Prokop, 2020). Accordingly, aberrations of MT bundles are sensitive indicators of axonal pathology, with disorganised curling of MTs being a widespread pathological phenotype (Fig.1; Hahn et al., 2019; Prokop, 2021; Smith et al., 2023).

**Fig. 1.**
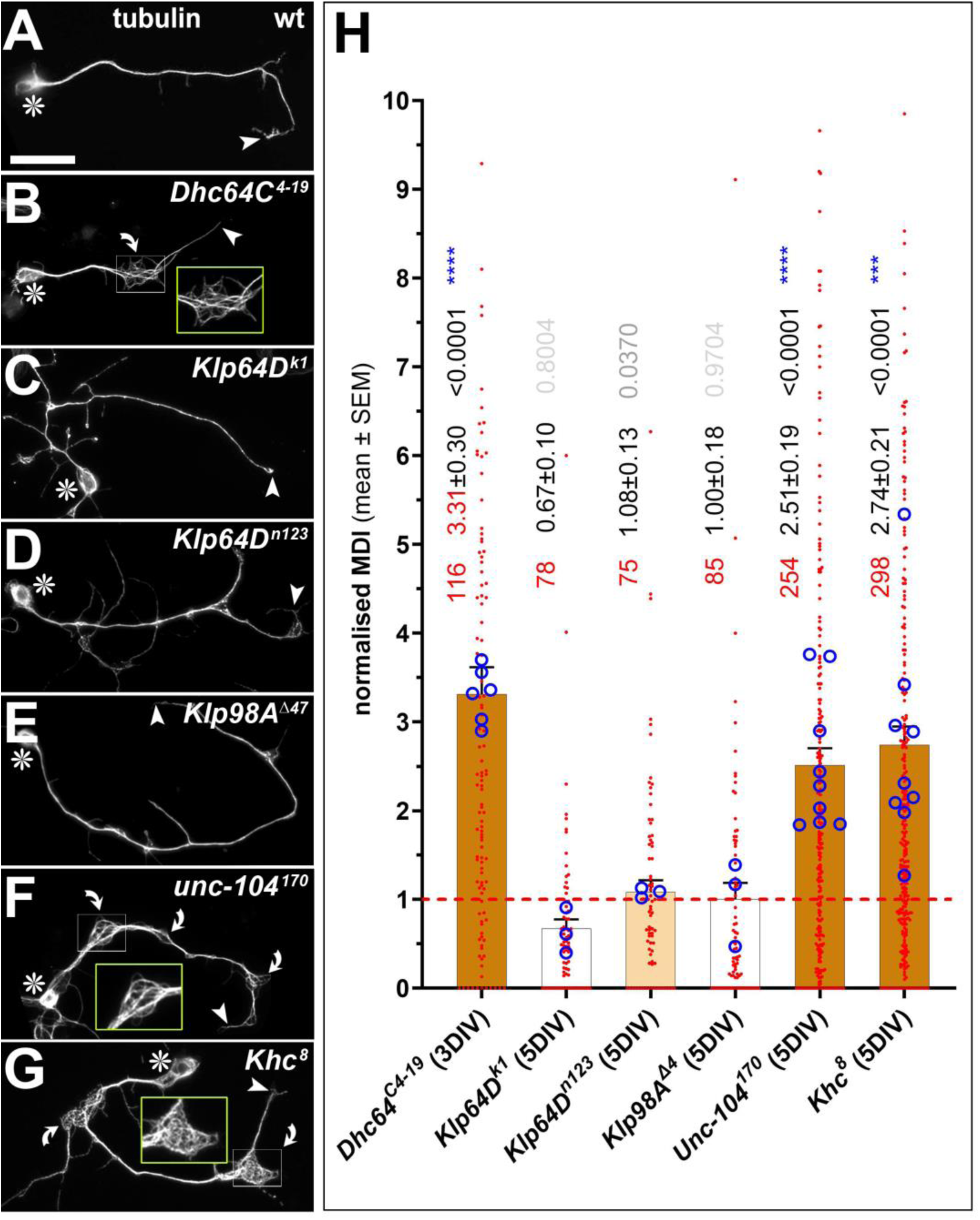
Deficiencies of three transport motor proteins cause MT curling. **A-G**) Examples of neurons displaying different mutations (as indicated top right) and stained for tubulin at 5DIV; asterisks indicate cell bodies, arrow heads axon tips, curved arrows areas of MT curling, white rectangles shown as twofold-magnified yellow-emboxed insets; scale bar in A represents 20µm in all images. **H**) Quantification of MT curling phenotypes measured as MT disorganisation index (MDI) and normalised to wild-type controls (red stippled line); mean ± SEM is indicated left of data points. Single data points are shown as dark-red dots and the means of replicates (independent coverslips from usually two experimental repeats) are shown as blue circles with their statistical significance (established using t-tests) indicated as blue asterisks; numbers of assessed neurons are shown in orange, their p values relative to controls (established by Mann-Whitney tests) in black or grey; the intensity of the brown bar colour reflects the degree of significance of the increased values.

Secondly, we used *Drosophila* primary neurons as a cost-effective model system, where mechanistic complexity can be addressed with fast and efficient genetic strategies (Hahn et al., 2021; Prokop et al., 2013; Qu et al., 2017). Loss-of-function analyses in *Drosophila* are facilitated by the fact that key factors, such as Kinesin heavy chain (Khc, kinesin-1), Kinesin light chain (Klc) or Milton, are each encoded by a single gene in *Drosophila*, as opposed to three, four or two mammalian orthologues, respectively. Furthermore, genetic tools are readily available to manipulate virtually any gene in question - and all these functional approaches can be combined with efficient and well-established readouts reflecting axonal physiology (Hahn et al., 2016; Prokop et al., 2013; Prokop et al., 2012; Sánchez-Soriano et al., 2010).

Here we show that the losses of three motor proteins, the KIF5 homologue Kinesin heavy chain (Khc; kinesin-1), the Kif1A/Bβ homologue Unc-104 (PH domain-containing kinesin-3), and the DYNC1H1 homologue Dynein heavy chain 64C (Dhc64C), all cause MT-curling and reduced numbers of mitochondria and synaptic dots in axons. Refined genetic analyses of Khc and its various adaptor proteins revealed two very different mechanisms that cause MT-curling: Firstly, loss of Khc-mediated transport of mitochondria and lysosomes causes harmful reactive oxygen species (ROS) which, in turn, induce MT-curling in fly and mouse neurons alike. Secondly, loss of the adaptor protein Kinesin light chain (Klc) causes overactivation of Khc as a ROS-independent curl-inducing mechanism mimicked by hyperactivating mutations of Khc; MT curling in these conditions occurs likely through direct force-mediated damage. Further experiments suggest that these fundamental mechanisms apply to other transport motor proteins. We discuss a conceptual model that can explain our findings and provide new explanations for observed phenomena in the neurodegeneration field.

## Results

### Losses of Khc, Unc-104 or Dhc cause axonal microtubule curling

To start our investigation, we assessed whether functional loss of motor proteins would cause measurable changes of axonal microtubule (MT) bundles as well-established indicators of axonal health (Okenve-Ramos et al., 2024; Prokop, 2020; Smith et al., 2023). We assessed MT bundle organisation in axons of neurons deficient for a range of transport motor proteins: (a) Dynein heavy chain (Dhc) is the obligatory motor-bearing subunit of the dynein/Dynactin complex essential for most, if not all, MT-based retrograde transport (Reck-Peterson et al., 2018); (b) Klp64D is one of two obligatory subunits of heterodimeric kinesin-2 (KIF3 homologue), reported to mediate anterograde axonal transport of acetylcholine-related synaptic enzymes or olfactory receptors (Baqri et al., 2006; Jana et al., 2021; Kulkarni et al., 2017; Ray et al., 1999); (c) the PX-domain-containing type 3 kinesin Klp98A (KIF16B homologue) was shown to mediate autophagosome-lysosome dynamics and endosomal Wingless transport in non-neuronal cells but is also strongly expressed in the nervous system (Mauvezin et al., 2016; Witte et al., 2020; flybase.org: <u>FBgn0004387</u>); (d) the PH-domain-containing type 3 kinesin Unc-104 (Kif1A homologue) is essential for synaptic transport in *Drosophila* axons (Pack-Chung et al., 2007; Voelzmann et al., 2016); (e) Kinesin heavy chain/Khc is the sole kinesin-1 in *Drosophila* (KIF5A-C homologue) involved in multiple transport functions in *Drosophila* neurons (see details below; e.g. Bowman et al., 2000; Gindhart et al., 2003; Glater et al., 2006; Loiseau et al., 2010; Rosa-Ferreira et al., 2018; Saxton et al., 1991).

We cultured primary neurons obtained from embryos homozygous for loss-of-function mutant alleles of these motor proteins (see Methods) and analysed them at 5DIV (days *in vitro*). These analyses revealed that especially the losses of Dhc, Khc and Unc-104 caused enhanced MT bundle disintegration displaying as areas where MTs are disorganised into curled, criss-crossing arrangements (from now on referred to as MT curling; curved arrows and enlarged insets in Fig.1,B,F-H).

To assess whether MT curling phenotypes were accompanied by transport defects, we analysed additional sub-cellular markers in *Khc^8^*, *unc-104^170^* and *Dhc64C^4-19^* homozygous mutant neurons. First, using the pre-synaptic protein Synaptotagmin (Syt) as a crude but easy to assess indicator of vesicular transport (Voelzmann et al., 2016), we found reduced presynaptic spots within axons of neurons mutant for any one of these three motor proteins (Fig.2A-D,P). Second, also the axonal number and distribution of mitochondria (visualised with mitoTracker; Klionsky et al., 2012), was significantly reduced in neurons lacking either Khc, Unc-104 or Dhc64C function (Fig.2H-K,O). Third, in *Khc^8^*mutant neurons, we also assessed the distribution of endoplasmic reticulum (ER) using a genomically GFP-tagged allele of Rtnl1, a constituent of axonal smooth ER (O’Sullivan et al., 2012). In wild-type neurons, ER is distributed evenly along the entirety of the axon (inset in Fig.S1A); loss of Khc does not affect this continuity, but about three quarters of neurons show an abnormal prominent accumulation of Rtnl-1::GFP label at their tips (see details and additional data in Fig.S1).

**Fig. 2.**
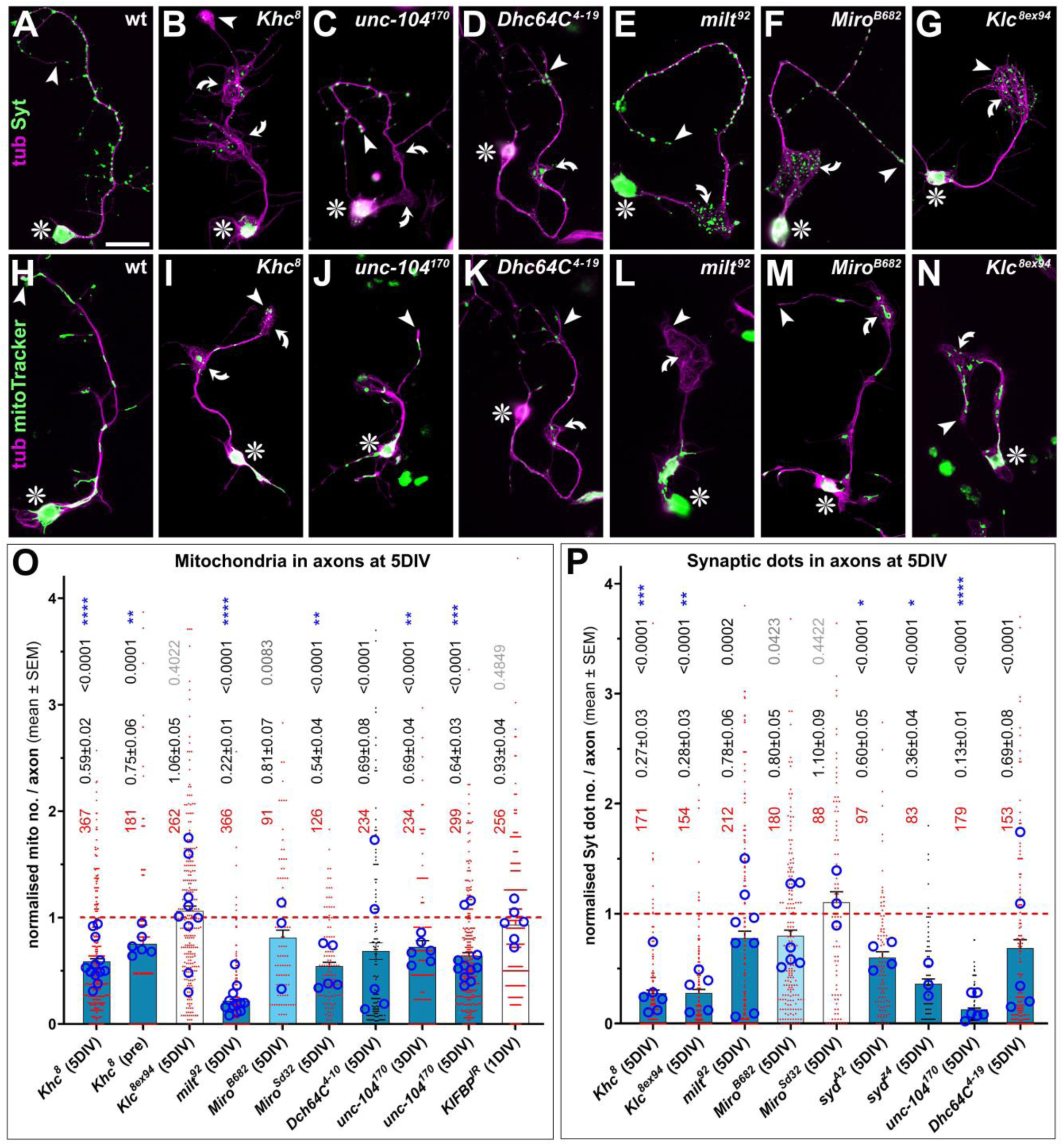
Impacts of motor protein and linker mutations on numbers of axonal mitochondria and Synaptotagmin-labelled spots. **A-N**) Examples of neurons displaying different mutations (indicated top right) and stained at 5DIV for tubulin (tub, magenta) and either Synaptotagmin (Syt, green in A-G) or with mitoTracker (green in H-N); scale bar in A represents 20µm in all images. **O,P**) Quantification of axonal numbers of Syt-positive spots (O) or mitochondria (P), all normalised to wild-type controls (red stippled line). Graphs are organised as indicated in Fig.1, only that the degree of significance of decreased values is indicated by blue bar colours of different shades.

In conclusion, our data strongly suggest that the loss of at least three motor proteins with demonstrated roles axonal transport cause MT curling.

### Khc displays strong maternal effects

We next focussed analyses on Khc, because many independent genetic tools are available enabling us to dissect sub-functions of Khc’s transport roles (Fig.3B). As the first step, we validated Khc’s MT-curling phenotype using the additional mutant allele *Khc^27^*, crossing *Khc^8^* and *Khc^27^* over a deficiency, and via RNAi-mediated knockdown of Khc (using pan-neuronal *elav-Gal4)*. In all cases, we found strong MT curling phenotypes at 5DIV confirming our initial finding (Fig.S2A,B).

**Fig. 3.**
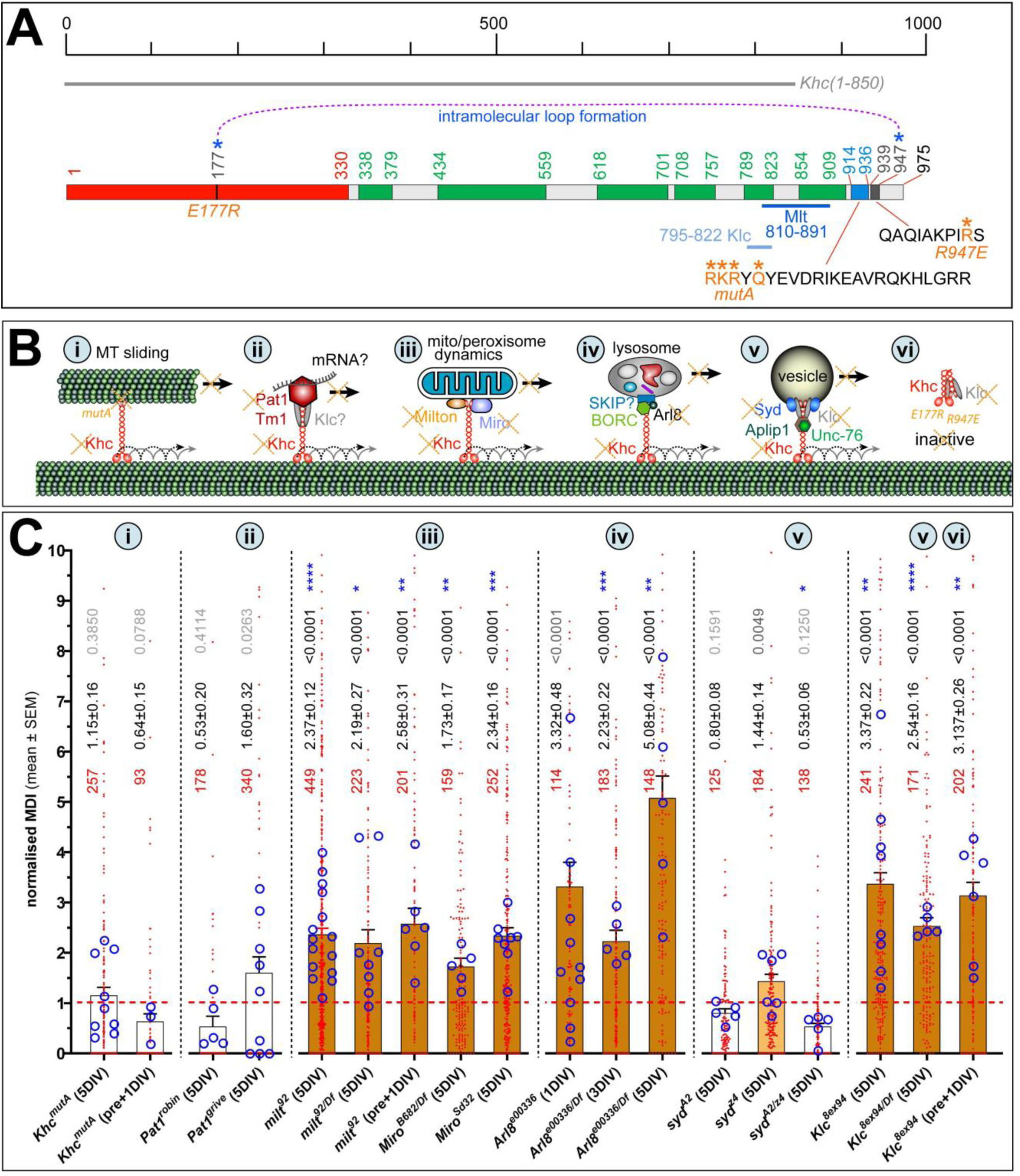
Khc subfunction deficiencies that cause MT-curling. **A**) Schematic representation of *Drosophila* Khc drawn to scale. Domains are colour-coded and start/end residues are indicated by numbers: motor domain (red; according to Sablin et al., 1996), coiled-coil domains required for homo- and/or heterodimerisation (green; as predicted by Ncoils in ensembl.org), the C-terminal ATP-independent MT-binding motif (blue; according to Winding et al., 2016), and the C-terminal auto-inactivation domain (dark grey; according to Kaan et al., 2011); the grey line above the protein scheme indicates the three expression constructs used in this study; below the protein scheme further details are shown: the sequence of the C-terminal MT-binding domain (*mutA* mutations indicated in orange; Winding et al., 2016), the sequence of the auto-inactivation domain (indicating the IAK motif and R947E mutation; Kelliher et al., 2018), the binding areas (darker green coiled-coils) of Klc (according to Veeranan-Karmegam et al., 2016) and Milt (known to overlap with Klc; Glater et al., 2006; Verhey et al., 1998). **B**) Schematic representation of some sub-functions of Khc (details and abbreviations in main text; red and stippled black lines indicate processive transport): via a C-terminal MT-binding domain Khc can slide MTs (i); by associating with Pat1 (and potentially Klc) it is expected to transport non-vesicular cargoes including mRNA (ii); with Milt and Miro it mediates transport of mitochondria and potentially peroxisomes (iii); its interaction with Arl8 is required to drive the transport of lysosomes and its derivatives (iv); a protein complex containing Klc and Syd is required for vesicular transport (v); in the absence of such associations Khc is auto-inhibited and detaches from MTs assisted by Klc (vi); to interfere with these subfunctions in this study, different genes were genetically removed (orange crosses) or specific *Khc* mutant alleles used (italic orange text). **C**) Quantified effects on MT curling caused by specific mutations affecting Khc sub-functions (numbers in grey circles indicate which function in A is affected); for details of graph organisation refer to the legend of Fig.1.

However, the MT phenotypes were not evident at earlier stages as revealed by analyses of *Khc^8/Df^* mutant neurons at 6 hours *in vitro* (HIV) or 3DIV (Fig.S2B,D): either phenotypes accumulate gradually over time (as observed for loss of Efa6; Qu et al., 2019), or mutant phenotypes are masked by maternal *Khc* gene product deposited during oogenesis by the heterozygous mothers (as previously observed for Eb1; Alves-Silva et al., 2012; Prokop, 2013).

To distinguish between these two possibilities, we used a pre-culture technique where neurons are kept in centrifuge tubes for 5 days to deplete maternal gene product, before growing them in culture (Prokop et al., 2012; Sánchez-Soriano et al., 2010). *Khc^8/Df^*-mutant neurons pre-cultured in this way, displayed prominent MT curling already 12 hrs after plating (Fig.S2C,E), suggesting that Khc has a prominent maternal contribution that persists for more than 3 days. Similar observations were made when pre-culturing neurons homozygous for *Khc^8^* or the *Khc^1ts^*mutant allele (details in Fig.S2C). We also tested whether maternal contribution might influence our data for other readouts, such as mitochondrial numbers: mitochondria might be transported into axons during the first days of axon growth and remain there when analysed at 5DIV. This would explain the stark difference between Khc and Milt deficiency (Fig.2O). However, mitochondria numbers in pre-cultured neurons seemed even less depleted than in neurons at 5DIV (Fig.2O) arguing against our initial suspicion (see further details in the Discussion).

### MT sliding functions of Khc do not link to MT-curling

We next tested potential mechanisms of Khc that might cause curling when abolished. For example, Khc might contribute to MT bundle maintenance by sliding MTs to achieve their even distribution along axons. Khc’s MT-sliding function is made possible by its C-terminal MT-binding site which, together with the MT-binding motor domain, enables it to cross-link MTs and move them against each other (Fig.3Bi; Andrews et al., 1993; Jolly et al., 2010; Lu et al., 2013; Lu et al., 2015; Winding et al., 2016).

To test whether Khc-mediated sliding contributes to MT bundle regulation we used the lethal *Khc^mutA^* mutant allele that specifically abolishes the binding of the Khc C-terminus to MTs but not other cargoes (Fig.3A; Winding et al., 2016). *Khc^mutA^*-mutant neurons were cultured in different ways: either as embryo-derived neurons cultured for 1DIV after 5d pre-culture, or as embryo-derived neurons cultured for 5DIV without pre-culture. In both cases, these neurons failed to show enhanced MT curling (Fig.3Ci), suggesting that the *Khc* mutant phenotype is not caused by the loss of its MT sliding function.

### Losses of Milton, Miro and Arl8 cause MT-curling phenotypes

Next, we focussed on Khc’ organelle transport functions focussing on mitochondria, peroxisomes and lysosome-related membrane compartments. The axonal transport of mitochondria requires the linker protein Milton and its binding partner Miro (a small GTPase) which anchor to the C-terminus of Khc (Fig.3A,Biii; Harbauer, 2017; Misgeld and Schwarz, 2017; Sheng, 2017; Smith and Gallo, 2018; Stowers et al., 2002). When testing the loss-of-function mutant alleles *milt^92^*, *Miro^Sd32^* and *Miro^B682^*, all caused prominent increases in MT curling at 5DIV (Fig.3Ciii). This phenotype correlated with a reduction in axonal mitochondria numbers (Fig.2L,M,O), as was similarly reported from *in vivo* studies (Glater et al., 2006; Guo et al., 2005; López-Doménech et al., 2018; Russo et al., 2009; Vagnoni et al., 2016).

At first sight, this suggested that defective mitochondrial transport can trigger MT curling. However, Miro has also been reported to mediate the transport of peroxisomes (Castro et al., 2018; Covill-Cooke et al., 2020; Okumoto et al., 2018; Tang, 2018). But a series of experiments convinced us that peroxisomes are negligible in the context of MT curling. Firstly, we labelled peroxisomes with PTS1::YFP (Faust et al., 2014) and found them mainly restricted to neuronal cell bodies (on average >10 peroxisomes) with axons displaying only an average of two (Fig.S3A-D). This is consistent with *in vivo* studies reporting high peroxisome abundance in glial cells but not neurons (Chung et al., 2020). Secondly, the culture medium we used contains only trace amounts of fatty acids (Else, 2020; Prokop et al., 2012; Schneider, 1964), so that peroxisomes appear negligible contributors to energy metabolism needed in only low numbers (Wanders et al., 2020). Thirdly, preliminary analyses suggested that loss of Khc has little impact on peroxisome distribution in axons (see details in Fig.S3E), and loss of peroxisomes (deletion of the biogenesis factor Pex3) did not at all increase MT curling (Fig.S3F).

In contrast to peroxisomes, lysosome-related membrane compartments are highly relevant. Khc-mediated lysosome transport requires Arl8 (ARL8A/B homologue; a small GTPase of the Arf family) in conjunction with SKIP/PLECHM2 (potential orthologue of Prd1 or Plekhm1 in fly) and the BORC complex (present in *Drosophila*; flybase.org; Fig. 3Biv; Keren-Kaplan and Bonifacino, 2021; Rosa-Ferreira and Munro, 2011; Rosa-Ferreira et al., 2018). Using a loss-of-function mutant allele of Arl8 in homozygosis or over deficiency, we observed strong MT curling at 5DIV (Fig. 3Civ).

Taken together, our results suggest that MT-curling observed in Khc-deficient neurons is, at least in part, mediated by aberrant transport of mitochondria and lysosomes, but not peroxisomes. Since Arl8 and TRAK/Milton were also implemented in retrograde transport (Canty et al., 2023; Paumier and Gowrishankar, 2024), they might contribute also to MT-curling observed upon loss of Dynein (Fig.1B,H).

### Excessive ROS triggers MT-curling in fly and mouse neurons

Reduced numbers of mitochondria or their wrong localisations are likely to impair important aspects of local axon homeostasis. Expected changes might include the regulation of calcium, ATP, Krebs cycle-derived metabolites, protein quality control, and signalling including reactive oxygen species (ROS; Glynn, 2017; Misgeld and Schwarz, 2017; Paupe and Prudent, 2018). Of these, energy depletion and ROS were shown to affect MT networks (Conze et al., 2024; Park et al., 2013; Shields et al., 2024). Since both mitochondrial and lysosomal dysfunction have been linked to oxidative stress (Kurz et al., 2008; Zhang et al., 2016), we tested ROS dyshomeostasis as a potential common nominator leading to MT curling.

For this, we first applied DEM (diethyl maleate) which causes ROS increase by inhibiting the anti-oxidant compound glutathione (‘blue arrowhead’ in Fig.4A; Albano et al., 2015; Dasgupta et al., 2012; Pompella et al., 2003). When applying 100 µM DEM for 12hrs before fixation (from 4.5 to 5DIV) we observed robust MT curling (Fig.4F). To validate these findings, we genetically manipulated the function of ROS-regulating enzymes (‘red arrowheads’ in Fig.4A); this included the use of a null allele of the peroxidase Catalase (*Cat^n1^*; Walker et al., 2018), loss and gain-of-function of Superoxide dismutase 1 (converting O₂^-^ into H_2_O_2_; *Sod1^n1^*, *Sod1^n64^*, *elav>Sod1*; Palma et al., 2020a), targeted expression of *Nox* (NADPH oxidase; enhancing O₂^-^ levels), of *Sod1* (to reduce O₂^-^ levels and enhance H_2_O_2_) and of *Duox* (Dual oxidase; to increase H_2_O_2_ levels; Anh et al., 2011; Bedard and Krause, 2007; Zelko et al., 2002). All these manipulations caused increased MT curling (Fig.4F). Of these, the observed effects upon loss of Catalase might be surprising when considering that peroxisomes (as the primary sites of localisation) are in low abundance; but Catalase was reported to enrich in the cytoplasm when peroxisomes are absent (Kleinecke et al., 2017; Zhou and Kang, 2000).

**Fig. 4.**
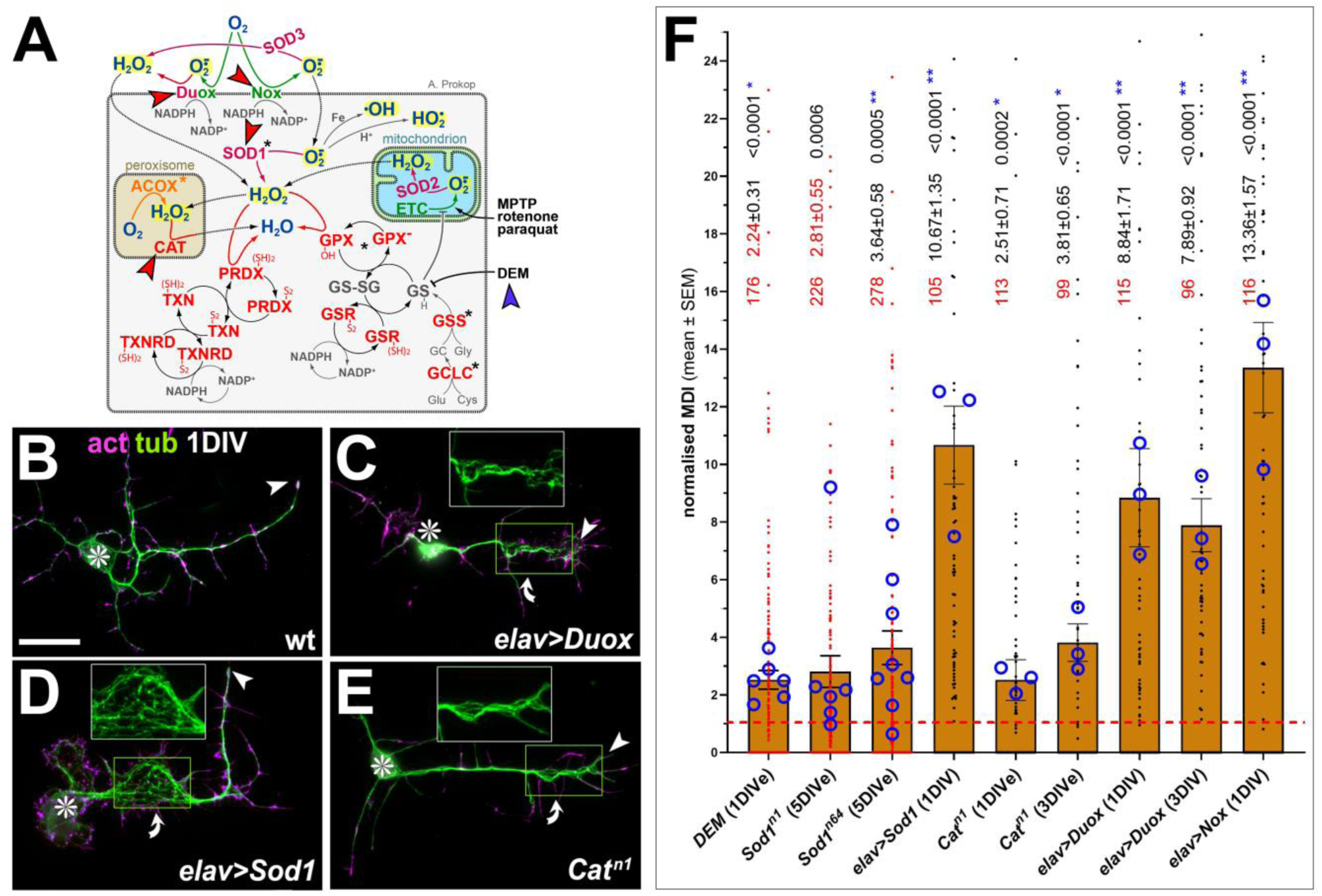
ROS enhancing manipulations cause MT curling phenotypes. **A**) Scheme illustrating the complexity of ROS-regulating systems (for molecular details refer to Smith et al., 2023); reactive oxygen species are highlighted in yellow, manipulated enzymes are indicated by red arrow heads and ROS-inducing DEM/diethyl maleate with a blue arrowhead. **B-E**) Examples of neurons, either wild-type (wt), expressing Sod1 or Duox (driven by *elav-Gal4*), or homozygous for *Cat^n1^*, all cultured for 1DIV and stained for actin (act, magenta) and tubulin (tub; green); asterisks indicate cell bodies, arrow heads axon tips, curved arrows areas of MT curling; yellow emboxed areas are shown as 1.5-fold enlarged insets (green channel only); scale bar in B represents 20µm in B-E. **F**) Quantification of MT curling phenotypes measured as MT disorganisation index (MDI) and normalised to wild-type controls (red stippled line); for further explanations of graph organisation refer to the legend of Fig.1.

Finally, we tested whether the MT-curl-inducing effects are conserved in mammalian neurons. We therefore applied 100 or 200 µM DEM to cultured mouse primary cortical neurons and found a robust increase in MT-curling (Fig.5).

**Fig. 5.**
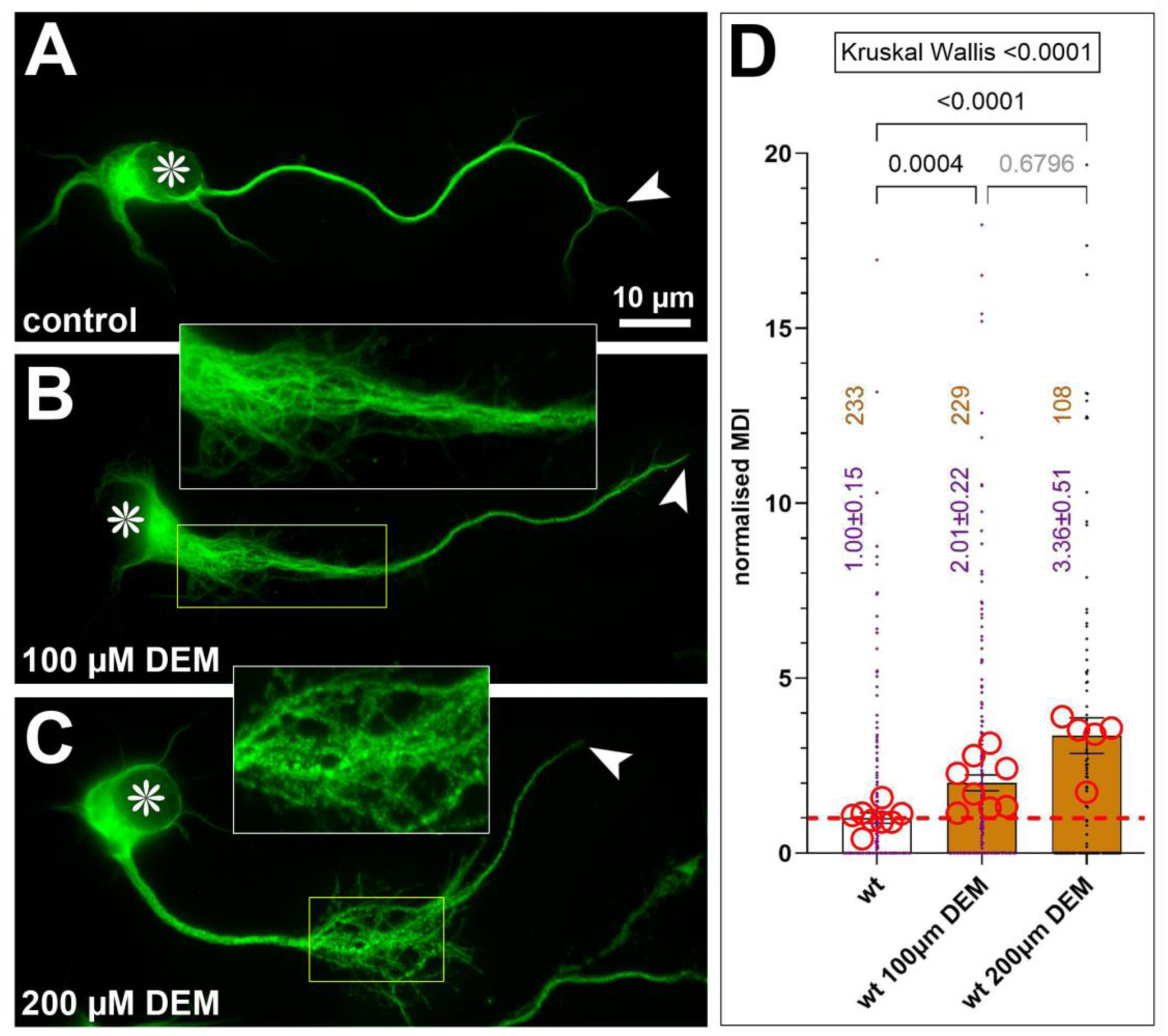
DEM induces MT-curling in primary mouse cortical neurons. **A-C)** Mouse primary neurons stained for tubulin, either untreated, or treated with 100µM or 200µM DEM, as indicated; asterisks indicate cell bodies, arrow heads axon tips; yellow boxes correspond to the areas shown as 2-fold magnified insets. **D)** Quantification of MT-curling data obtained. Significance was assessed by Kruskal Wallis test followed by Dunn’s multiple comparisons posthoc tests. For explanations of graph organisation, refer to the legend of Fig.1; red circles represent 5-8 independent coverslips per condition obtained from 3 independent mouse primary cortical neuron preparations.

Taken together, dyshomeostasis of ROS clearly triggers MT-curling in fly and mouse neurons alike. Upregulation of either O₂^-^ or H_2_O_2_ seems to cause this effect, although O₂^-^ is perhaps more likely to elicit its effects through conversion into H_2_O_2_ (Bedard and Krause, 2007; Palma et al., 2020b).

### MT-curling downstream of Khc, Milt and Arl8 deficiency seems mediated by harmful ROS

To assess whether harmful ROS might be responsible for linking aberrant organelle transport to MT-curling, we treated Khc-, Milt- and Arl8-deficient neurons with Trolox (6-hydroxy-2,5,7,8-tetramethylchroman-2-carboxylic acid). Trolox is an α-tocopherol/vitamin E analogue that displays beneficial antioxidant effects in many cell systems, including neuronal models of neurodegeneration; it inhibits fatty acid peroxidation and quenches singlet oxygen and superoxide (Chow et al., 1994; Giordano et al., 2020; Janc and Müller, 2014). To first validate its effect, we treated neurons with DEM to induce MT-curling; we observed that this phenotype was abolished upon co-application of Trolox (Fig.6A; see also Shields et al., 2025).

**Fig. 6.**
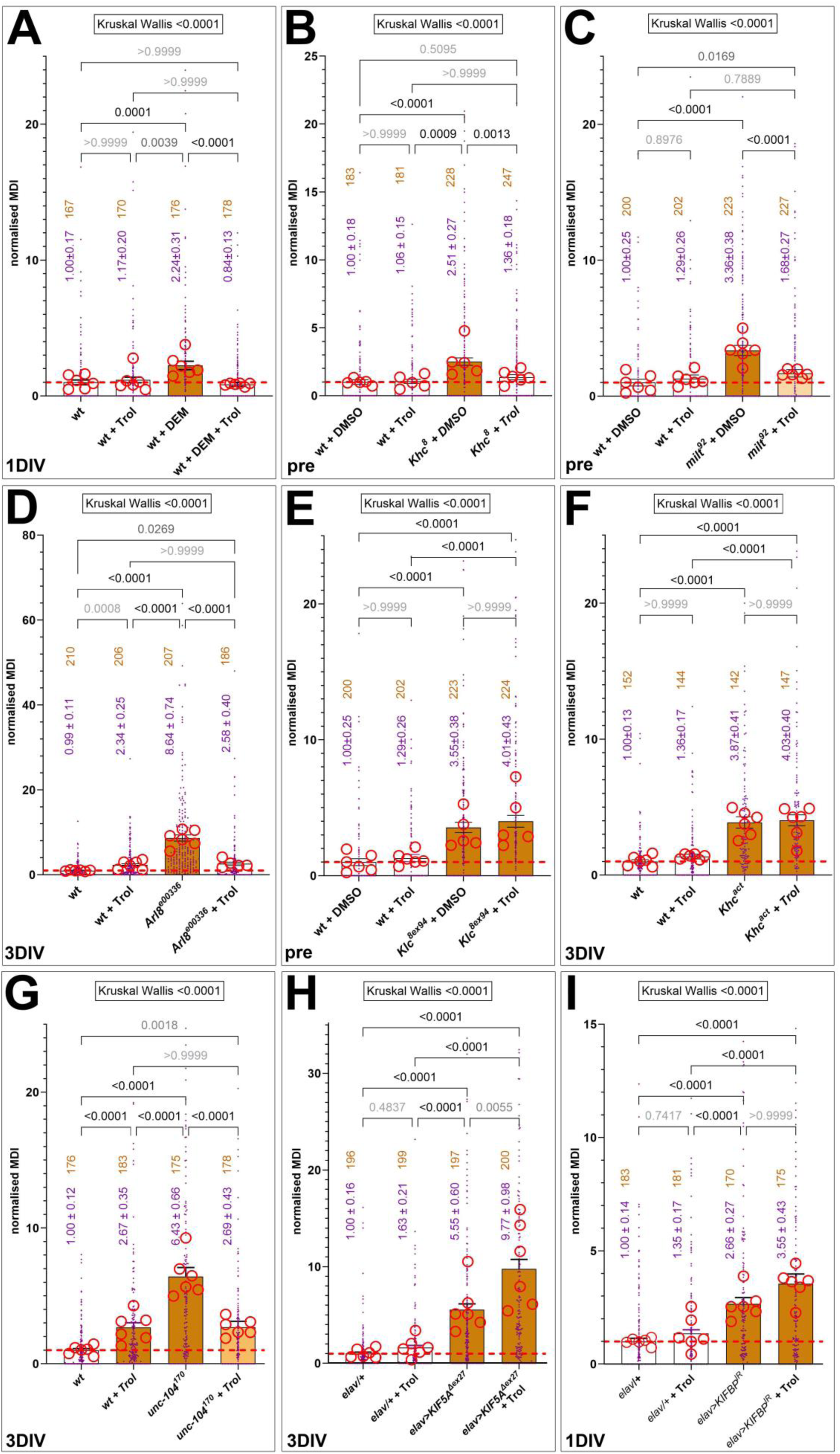
Ameliorating effects of Trolox on mutant MT-curling phenotypes. Each graph shows the results of separate experiments quantifying MT-curling phenotypes measured as MT disorganisation index (MDI) and normalised to wild-type controls (red stippled line); in each experiment, wild-type neurons with or without Trolox and mutant or DEM-treated neurons with or without Trolox treatment are shown; further culture conditions are indicated: which are either pre-culture (pre), or culture for 1 or 3DIV. Significance was assessed by Kruskal Wallis test followed by Dunn’s multiple comparisons posthoc tests. Further explanations of graph organisation can be taken from the legend of Fig.1.

We next treated *Khc^8^*, *milt^92^* and *Arl8^e00336^* mutant neurons with 100µM Trolox and found a robust suppression of the MT-curling phenotype in all three conditions (Fig.6B-D), suggesting that absence of their function causes harmful ROS as a key mediator of their MT-curling phenotypes (Fig.S5B,C). This also agrees with reports that rat neurons depleted of the Khc homologue KIF5C are more vulnerable to H_2_O_2_ application (Iworima et al., 2016).

### Klc causes MT curling independent of its role in vesicular transport

Also the transport of non-organelle cargoes, such as RNA-binding proteins, has been linked to axon degeneration (Feng et al., 2025) and might therefore cause MT-curling. For example, Pat1 (Protein interacting with APP tail-1) was shown to link Khc to non-vesicular transport in *Drosophila* oocytes (Fig.3Bii; Loiseau et al., 2010); it is also strongly expressed in the *Drosophila* nervous system ((flybase.org: <u>FBgn0029878</u>) although its potential neuronal cargoes are unknown. However, when assessing Pat1-deficient primary neurons, no convincing MT-curling was observed (Fig.3Cii).

We further tested loss of Khc-mediated vesicular cargo transport which requires a complex of various factors including Kinesin light chain (Klc; homologue of KLC1-4) and Sunday driver (Syd, a JNK-Interacting Protein; Fig.3Bv; Gindhart et al., 2003; Horiuchi et al., 2005; Koushika, 2008). Functional losses of Klc, Syd or Khc were shown to affect vesicular transport in larval axons to similar degrees, suggestive of mutual requirements (Bowman et al., 2000; Füger et al., 2012; Gauger and Goldstein, 1993; Gindhart et al., 1998; Hurd and Saxton, 1996; Pilling et al., 2006). Using primary neurons, we found that the number of synaptic Syt-labelled dots was reduced upon the losses of Khc, Klc and Syd and far less upon loss of Milt or Miro, whereas loss of Unc-104 showed the strongest depletion (Fig.2P). These data agree with the existence of Khc/Klc/Syd-mediated vesicular transport in *Drosophila* primary neurons, in parallel to known functions of Unc-104 in synaptic transport (Pack-Chung et al., 2007; Voelzmann et al., 2016).

However, despite their very similar inhibitory effects on vesicular transport, neurons lacking Syd showed no convincing increase in MT-curling, whereas loss of Klc caused a very strong phenotype (Fig.3Cv). These findings might suggest that MT-curl-inducing effects upon loss of Klc do not relate to vesicular transport. This notion was corroborated by knock-down of GAPDH (glyceraldehyde-3-phosphate dehydrogenase), an enzyme required for glycolysis; its loss in *Drosophila* was previously shown to inhibit the axonal transport of vesicles but not of mitochondria (details in Fig.S4A,B; Zala et al., 2013). Consistent with these reports, knock-down of GAPDH in primary *Drosophila* neurons caused robust reduction of synaptic dots and a far more moderate effect on mitochondrial numbers (Fig.S4C,D). However, GAPDH knock-down did not cause any increase in MT curling phenotypes (Fig.S4C), further supporting our conclusion that reduced vesicular transport is not a MT-curl-inducing condition.

To explore potential alternative mechanisms underlying the Klc-deficient MT-curling phenotype, we tested the potential involvement of ROS. However, when applying Trolox, the MT-curling phenotype in Klc-deficient neurons was not at all suppressed (Fig.6E), clearly differing from loss of Khc, Milt or Arl8 (Fig.6B-D). This finding further suggested the involvement of a very different mechanism.

### Excessive pools of active Khc might explain the Klc-deficient MT-curling phenotype

To gain a mechanistic understanding of Klc’s role in MT regulation, we first performed genetic interaction studies. These studies assessed the potential occurrence of MT-curling when combining heterozygous conditions of independent genes in the same animals or cells (referred to as trans-heterozygous condition); phenotypes occurring upon trans-heterozygosis are suggestive of a direct or indirect functional interaction between the tested genes. Unfortunately, the genetic interactions we tested were not robust enough to draw clear conclusions, but they showed meaningful trends. When combining *Klc^8ex94/+^* with *milt^92/+^*, the trans-heterozygous condition does not exceed the sum of the single heterozygous conditions; this might suggest absence of genetic interaction in agreement with their potential mechanistic dichotomy (Fig.S6A). In contrast, the combination of *Khc^8/+^* and *milt^92/+^* caused a phenotype that seemed stronger than the sum of the single heterozygous conditions (Fig.S6B), mildly suggestive of genetic interaction and agreeing with their common function in mitochondrial transport. Finally, *Khc^8/+^* and *Klc^8ex94/+^* showed a trend towards a negative interaction (Fig.S6C), as if Khc acts as a mediator of the Klc-deficient phenotype.

The latter finding might point at reported roles of Klc in the inactivation of Khc: Khc molecules not engaged in transport transit into a labile auto-inhibited and MT-detached Λ configuration where the C-terminus binds intramolecularly to the N-terminus (‘stippled blue line’ in Fig.3A); Klc promotes this inactive stage (Figs.3Bvi; Bowman et al., 2000; Chiba et al., 2022; Koushika, 2008; Smith et al., 2024; Verhey and Hammond, 2009; Verhey et al., 1998; Weijman et al., 2022; Wong and Rice, 2010; Yip et al., 2016). In non-neuronal cells, overriding auto-inhibition of the Khc-Klc complex causes severe MT-curling (Paul et al., 2020; Randall et al., 2017).

To test whether Klc-deficient MT-curling in neurons might involve pools of hyperactivated Khc (Fig.S5D), we first targeted the expression of GFP-tagged Khc constructs to neurons. Full-length Khc^FL^::GFP was homogeneously distributed along axons as similarly reported for its homologue KIF5A (Cozzi et al., 2024); it did not cause a MT-curling phenotype at 5DIV (Fig.7A,C). Next, we expressed the deletion construct Khc^1-850^::GFP (indicated as grey bar in Fig.3A) which lacks the C-terminal region required for auto-inhibition, cargo transport and MT sliding. Khc^1-850^::GFP accumulated at axon tips (Fig.7B), as is typical of non-inactivating kinesins (Nakata and Hirokawa, 2003; Niwa et al., 2013; Seiler et al., 2000); it caused a modest MT-curling phenotype (Fig.7D). These results might suggest that free-running, non-inactivating Khc can harm axonal MT bundles, unlike Khc^FL^ which inactivates and detaches from MTs when unengaged with cargo (Fig.S5F,G).

**Fig. 7.**
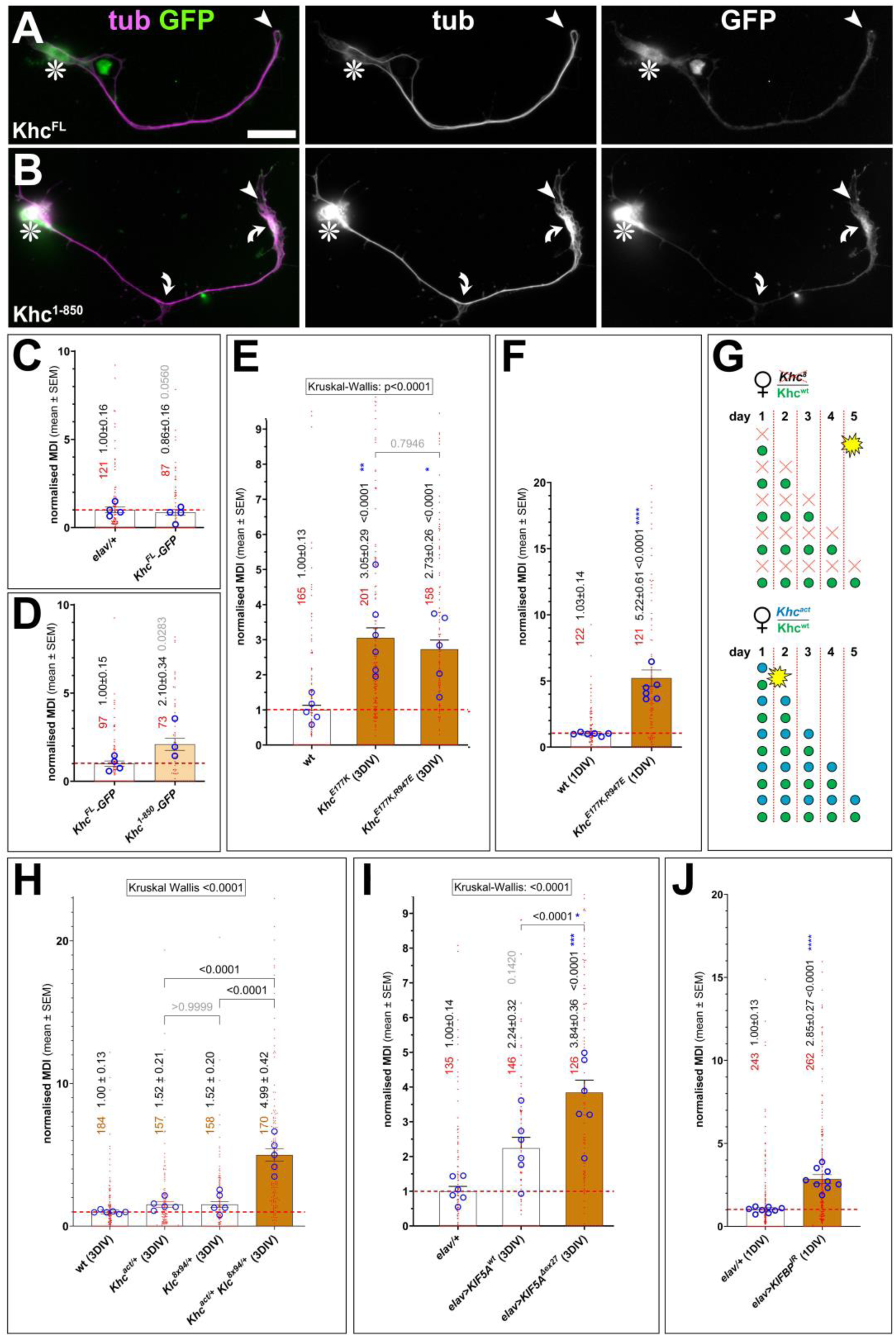
Impacts of activated Khc on MT curling. **A,B**) Examples of neurons at 5DIV expressing different Khc constructs (indicated bottom left; compare Fig.3A) and stained for tubulin (tub, magenta) and GFP (green), also shown as greyscale single channel images on the right; asterisks indicate cell bodies, arrow heads axon tips, curved arrows areas of MT curling; scale bar in A represents 20µm in all images. **C-F,H-J**) Quantification of MT curling phenotypes measured as MT disorganisation index (MDI) and normalised to wild-type controls (red stippled lines); genotypes are shown below, also indicating the culture period (5DIV, 3DIV). Significance of single comparisons was assessed by Mann Whitney tests of multiple comparisons by Kruskal Wallis test followed by Dunn’s multiple comparisons posthoc tests. For descriptions of graph organisation, refer to Fig.1. **G)** Illustration of the maternal Khc^wt^ product (green dots) deposited in the eggs by heterozygous mothers and decaying over ∼5 days; in *Khc^8^* mutant neurons, the phenotype (yellow star) appears upon maternal product depletion (top); in non-inactivating *Khc^act^* mutant neurons, the phenotype occurs early, suggesting that the presence of Khc^act^ protein (blue dots) overrides the healthy function of Khc^wt^.

Since Khc^1-850^::GFP cannot bind cargo, it likely imposes less force on MTs than a loaded kinesin. We therefore tested the genomically engineered *Khc^E177K^* and *Khc^E177R, R947E^* mutant variants which are non-inactivating but still able to transport cargo: the E177 and R947 residues are essential for the C- to N-terminal interaction (‘blue asterisks’ in Fig.3A; Kaan et al., 2011; Kelliher et al., 2018); their mutation suppresses auto-inhibition and causes lethality and distal accumulations of Khc, but it does not affect the ability to transport cargo (Brendza et al., 1999; Kelliher et al., 2018). When we cultured *Khc^E177K^* and *Khc^E177R, R947E^* homozygous mutant neurons, they displayed robust MT-curling when tested at 3DIV (Figs.7E, S5E). Surprisingly, this is at a stage at which Khc-deficient neurons do not show a phenotype yet (Fig.S2B), as became even clearer when testing *Khc^E177R, R947E^* mutant neurons at 1DIV which resulted in robust MT-curling (Fig.7F). This strongly suggests that the E177K and R947E point mutations cause dominant phenotypes, overriding the wild-type maternal protein present during the first days of development (Fig.7G).

Finally, we tested whether phenotypes observed upon Klc loss and Khc non-inactivation are functionally related. Firstly, as observed in Klc-deficient neurons, Trolox failed to suppress the *Khc^E177R, R947E^* mutant MT-curling phenotype (Fig.6E,F). Secondly, the trans-heterozygous combination of *Klc^8ex94/+^* with *Khc^E177R, R947E/+^* in primary neurons, revealed a very strong and convincing genetic interaction that supports synergetic contributions of Klc and intramolecular loop formation during Khc auto-inhibition (Fig.7H).

Taken together, we conclude that MT-curling observed upon loss of Klc is likely due to failed inactivation of Khc. This represents a very different pathological mechanism from that observed with losses of Khc, Milt, Miro or Arl8, although the MT-curling phenotype of all these conditions looks surprisingly similar.

### Loss and gain of other motor proteins likewise cause MT-curling

So far, our studies of the *Drosophila* kinesin-1 Khc suggest that MT-curling can be due to gain-of-function through failed auto-inhibition or caused by loss-of-function through harmful ROS due to impaired organelle transport. We next asked whether these two mechanistic pathways might be more widely applicable. For this, we assessed the PH domain-containing *Drosophila* kinesin-3 Unc-104 and the human kinesin-1 KIF5A.

As explained above, Unc-104 causes MT-curling (Fig.1F,H). Previous research in larval nerves failed to resolve whether Unc-104 transports mitochondria (Barkus et al., 2008). Our analyses in *unc-104^170^* mutant neurons at 3DIV and 5DIV show that axonal mitochondria numbers are reduced to similar degrees as observed upon loss of Khc (Fig.2I,J,O), suggesting a direct or indirect role in mitochondrial motility. We speculated therefore that Unc-104 might cause ROS dyshomeostasis. This notion was confirmed when applying Trolox, which robustly suppressed the MT-curling phenotype in *unc-104^170^* mutant neurons, as similarly observed for loss of Khc (Figs.6B,G).

Like kinesin-1, also kinesin-3 motors undergo inactivation involving intramolecular loop formation (Cong et al., 2021; Niwa et al., 2024). Unfortunately, the non-inactivating *unc-104^bris^* mutant allele described for *Drosophila* (Kern et al., 2013; Niwa et al., 2024) seems no longer to exist. As an alternative strategy to assess potential effects of hyperactivated kinesin-3 on MT-curling, we turned to KIF-binding protein (KIFBP). KIFBP was initially described as a linker for KIF1Bα (lacking a PH domain) mediating mitochondrial transport (Nangaku et al., 1994; Wozniak et al., 2005). However, upon knock-down of KIFBP in *Drosophila* primary neurons, we did not observe any reduction in axonal mitochondria numbers (Fig.2O). Alternatively, newer studies suggest KIFBP to be involved in the inactivation of several kinesin classes, among them all kinesin-3 paralogues (Kevenaar et al., 2016; Solon et al., 2021). Consistent with such a role, we found that knock-down of KIFBP caused robust MT curling in primary neurons at 1DIV (Fig.7J), and this phenotype was not reduced when applying Trolox (Fig.6I). These findings suggest that, like Klc deficiency, also the loss of KIFBP seems to cause MT-curling through the mechanical route likely involving hyperactivation of Unc-104.

Finally, we used overexpression of wild-type and mutant KIF5A: the KIF5A^Δexon27^ mutant isoform is an aberrant splicing product linked to ALS characterised by a small C-terminal truncation substituted by a 39 amino acid-long neopeptide (orange in Fig.S6I); this weakens its ability to auto-inactivate, it promotes aberrant protein interactions trapping other mutant or wild-type KIF5A molecules, and causes a decrease of protein lifetime in a proteasome-dependent manner (Fig.S5I; Baron et al., 2022; Carrington et al., 2024; Cozzi et al., 2024; Nakano et al., 2022; Pant et al., 2022; Pino et al., 2023). Its overexpression might therefore cause curling either through gain-of-function by being over-activated, or loss-of-function by trapping endogenous kinesins.

We started our KIF5A investigation by expressing a C-terminally GFP-tagged wild-type version (KIF5A^wt^::GFP; Pant et al., 2022). GFP fluorescence was above detection level in 67% of cells but often occurring in aggregates, with 83% of fluorescent cells showing accumulations at axon tips (Fig.S7A,A’). Tip localisation suggested reduced auto-inhibition, as was further supported by a (non-significant) bias to cause MT-curling (Fig.7I). Comparable results were obtained with the Khc^1-850^::GFP construct (Fig.7D), suggesting that KIF5A might be hyperactivated but fail to bind to cargo linkers (Fig.S5G,H), despite the fact that the cargo linker-binding regions of KIF5A and Khc appear reasonably well-conserved (Fig.S8).

Kif5A^Δex27^::GFP also occurred in aggregates, with over 70% of fluorescent neurons showing tip localisation as a potential indicator of over-activation (FigS7B,C). However, the number of neurons displaying GFP above detection levels was below 50% when expressing Kif5A^Δex27^::GFP, as compared to 70% for KIF5A^wt^::GFP. This reduction might reflect the previously reported shorter protein lifetime of Kif5A^Δex27^::GFP mentioned above (Baron et al., 2022; Cozzi et al., 2024; Pino et al., 2023). Despite this reduction, expression of the mutant isoform displayed strong and robust MT-curling in *Drosophila* primary neurons at 3DIV (Fig.6I). To test whether this phenotype is due to over-activation of KIF5A, or to loss-of-function by trapping endogenous motors, we applied Trolox. We found that blocking ROS did not only fail to suppress the curling phenotype, but curling was consistently and robustly increased (Fig.6H). Increased MT-curling correlated with the observation that also the number of neurons with detectable GFP was increased from ∼50% without Trolox to ∼70% with Trolox, potentially suggesting a positive role of ROS signalling in Kif5A^Δex27^ degradation. Preliminary experiments with HEK293T cells expressing KIF5A^Δex27^::GFP showed a similar trend to increased protein levels when treated with Trolox (Fig.S9) potentially suggesting a conserved mechanism.

In conclusion, the experiments with Unc-104, KIFBP and KIF5A suggest that MT-curling upon loss- and gain-of-function is not specific to Khc, but is likely to apply also to other transport motors.

## Discussion

### A standardised *Drosophila* primary neuron system reveals potential patho-mechanisms linked to axonal transport

Transport motors and their linker proteins mediate the correct delivery and relocation of molecules and organelles required to maintain axonal physiology and function, yet their links to axonopathies remain poorly understood and speculative (Coleman, 2005; Guo et al., 2020; Kawaguchi, 2013; Sleigh et al., 2019). True understanding of axonal transport is confounded by various factors, such as the parallel or even interdependent involvement of different motor protein classes, or the involvement of many different cargo adaptor proteins that may interact promiscuously with different cargos and/or motors (Cason and Holzbaur, 2022; Hancock, 2014; Hirokawa et al., 2010).

Here we embraced this complexity using one standardised *Drosophila* primary neuron system amenable to efficient combinatorial genetics, combined with MT-curling as an efficient readout. Previous studies had already shown that *Drosophila* primary neurons provide powerful means to overcome redundancy or the complexity of genetic networks (e.g. Beaven et al., 2015; Gonçalves-Pimentel et al., 2011; Hahn et al., 2021; Qu et al., 2017). Here we used this system to investigate how aberrant axonal transport may cause neurodegeneration. This involved analyses of 30 different genetic loss- or gain-of-function conditions covering 18 different genes, partly in combination or applying different experimental regimes, and further complemented with DEM and Trolox treatments. Our loss-of-function studies were facilitated by the lower genetic redundancy as compared to vertebrates (e.g. Khc/KIF5: 1 gene in fly *vs.* 3 paralogues in mammals; Miro/RHOT: 1 *vs.* 2; Milt/TRAK: 1 *vs.* 2; Klc: 1 *vs.* 4; Unc-104/Kif1: 1 *vs.* 2; Arl8 1 *vs.* 2) rendering the need of double- or triple-mutant approaches obsolete.

In our view, MT-curling is not only an easy-to-quantify feature conserved between fly and vertebrate axons (Fig.5; Ahmad et al., 2006; Hahn et al., 2021; Rafiq et al., 2022; Sánchez-Soriano et al., 2009; Voelzmann et al., 2024), but it is a good indicator of lost axon integrity: MT bundles form the life-sustaining transport highways to communicate with the neuronal cell body and properly localise organelles and materials, thus sustaining cell biological processes. Accordingly, MT-curling affects axonal transport and negatively impacts axon health (Okenve-Ramos et al., 2024; Prokop, 2020; Smith et al., 2023).

Using MT-curling as a readout upon genetic and pharmacological manipulations in the standardised *Drosophila* primary neuron system, we found that phenotypes upon loss of Khc, Unc-104 and even Dhc are comparable with respect to enhanced MT curling and reduced numbers of mitochondria and synaptic dots in axons (Figs.1, 2). This functional overlap agrees with reports that these three motors might collaborate during transport or even display a degree of mutual dependency (Arpağ et al., 2019; Cason and Holzbaur, 2022; Zahavi et al., 2021). Therefore, the three motors might link to axonopathy through comparable mechanisms. Our data support this notion (further details below) and suggest two principal mechanisms underpinning MT-curling (Figs.8, S5): (1) motor loss disturbs axonal homeostasis leading to physiological imbalance and MT-curl-inducing ROS production (Figs.4 and 6A-D,G), whereas (2) hyperactivation causes ROS-independent mechanical imbalance where motor forces overwhelm the MT bundle maintenance machinery (Figs.6E,F,H,I and 7).

### Organelles regulate ROS homeostasis required for MT bundle maintenance

Our finding that harmful ROS is a key trigger of MT curling is consistent with reports in other systems: ROS can oxidise cysteins in both α- and β-tubulin (Landino et al., 2002; Löwe et al., 2001; Luduena and Roach, 1991) or lead to functional modifications of MT-regulating proteins such as Eb1, MAP2, MAP1B, tau or the collapsin response mediator protein 2 (CRMP2; Conze et al., 2024; Landino et al., 2004; Mattson et al., 1997; Miglietta et al., 1991; Shields et al., 2025; Sultana et al., 2013). ROS-induced modifications disrupt MT organisation in non-neuronal cells (Goldblum et al., 2021; Wilson et al., 2016; Wioland et al., 2021); they were reported (1) to inhibit axon growth and be causative in neurodegeneration (Neely et al., 1999; Wali et al., 2016), (2) to induce axon swellings in models of Parkinson’s disease, multiple sclerosis or motor neuron disease (Czaniecki et al., 2019; Nikić et al., 2011; Song et al., 2013) and (3) to block axonal transport and cause axonal MT disorganisation that could be reverted with ROS scavengers (Nikić et al., 2011; Shields et al., 2025; Sorbara et al., 2014).

Our data suggest that ROS-dependent MT-curling is caused by aberrant transport of mitochondria or lysosomes (Fig.3). In the case of lysosomes, this might be the indirect consequence of protein stress caused by failed protein removal, or their direct involvement in the prevention of oxidative stress (Paumier and Gowrishankar, 2024; Zhang et al., 2016). ROS production upon aberrant mitochondrial transport, is far less clear. For example, *milt* mutant neurons showed strong ROS-induced MT-curling in axons, although mitochondria frequently remain in cell bodies (Figs.2L,O and 3Ciii). Any excessive ROS generated by these somatic mitochondria would hardly reach far into the axon due to the presence of abundant ROS-buffering systems (Fundu et al., 2019; Kükürt et al., 2021; Oswald et al., 2018). Therefore, even the absence of mitochondria from axons might affect ROS homeostasis, suggesting they may play an active role in cytoplasmic ROS regulation (e.g. via Sod2-mediated H_2_O_2_ signalling; Palma et al., 2020b).

A further puzzling aspect of *milt* mutant neurons is the strong reduction in axonal mitochondria, which is far greater than upon loss of any motor protein, even when depleting Khc’s maternal component (Fig.2O). This might suggest Milt as a potential ’master regulator’ of mitochondrial transport in fly neurons. Milt is known to link to Khc and Dynein in both flies and mammals (Russo et al., 2009; van Spronsen et al., 2013), whereas no such reports exist for Unc-104 or vertebrate KIF1. So far, only KIF1Bα was suggested to perform mitochondrial transport via KIFBP (Nangaku et al., 1994; Wozniak et al., 2005). Therefore, our observed involvement of Unc-104 in mitochondrial transport and the ‘master regulator’ role of Milt remain puzzling. They may eventually find explanations through the above-mentioned intricate collaboration and interdependence of kinesin-1, kinesin-3 and dynein during axonal transport. The milder phenotype of Miro deficiency compared to Milton (Fig.2O) is consistent with findings in fly neurons *in vivo* as well as in mouse neurons (Glater et al., 2006; Guo et al., 2005; López-Doménech et al., 2018; Russo et al., 2009; Vagnoni et al., 2016; see also Mattedi et al., 2023); this may suggest that Miro is a facilitator but not a compulsory component of the mitochondria linker complex.

### Motor hyperactivation as a further factor leading to MT-curling

Our findings that motor hyperactivation causes MT-curling aligns well with published data: (1) *In vitro* experiments demonstrate kinesin-1-induced MT damage (Andreu-Carbó et al., 2021; Budaitis et al., 2021; Dumont et al., 2015; Triclin et al., 2021; VanDelinder et al., 2016). (2) MT curling is observed in *in vitro* gliding assays utilising kinesin-1 (Hahn et al., 2019; Lam et al., 2016; Nasirimarekani et al., 2022). (3) Kinesore-mediated activation of kinesin-1 in non-neuronal cells triggers dramatic MT-curling (Paul et al., 2020; Randall et al., 2017). (4) MT-curling in non-neuronal cells was suggested to be caused by anterograde transport (Bicek et al., 2009; Blob et al., 2025).

That hyperactive motors are potentially damaging, is demonstrated by the fact that neurons dispose them into glia cells (Xie et al., 2024) and by the fact that hyperactivating mutations of KIF5A and of *Drosophila* Khc cause ALS or lethality, respectively (Baron et al., 2022; Kaan et al., 2011; Kelliher et al., 2018; Pant et al., 2022; Pino et al., 2023). Furthermore, KLC2 has been linked to neuropathy (SPOAN; #<u>609541</u>) and KLC4 mutations have recently been reported to cause excessive axon branching (Haynes et al., 2022), which may be due to transport deficits or hyperactivation of kinesin-1. Similarly, non-inactivating mutations of Kif1A cause axonopathy (spastic paraplegia; Chiba et al., 2019; Gabrych et al., 2019), and KIFBP mutations impair axon growth, are disruptive to axonal MT bundles (Lyons et al., 2008) and cause the devastating neurological Goldberg-Shprintzen syndrome (GOSHS; Chang et al., 2019; Hirst et al., 2017) potentially involving kinesin hyperactivation as an underlying patho-mechanism.

## Conclusion

The key outcome of our studies is that one common phenotype of MT-curling (as an indicator of axon decay) can be generated through two distinct mechanisms: a physiological mechanism involving ROS dyshomeostasis upon motor protein or linker loss, or a mechanical mechanism due to hyperactivation of motor proteins. Our findings have essentially contributed to the formulation of the "dependency cycle of axon homeostasis" model (Prokop, 2021; Smith et al., 2023; details in Fig.8). It proposes that kinesins trailing along MT bundles during axonal transport (’2’ in Fig.8) pose a mechanical challenge that leads to MT-curling (’3’); this challenge is enhanced upon hyperactivation of motors (‘large red arrow’ in Fig.8). Active machinery of MT-regulating proteins and support through the cortical actin-spectrin sleeve is required to prevent disintegration and maintain axon bundles long-term (’4’). However, this machinery of MT bundle maintenance is itself dependent on materials and physiology provided by axonal transport (’5’); this aspect of the cycle is affected by loss of (organelle) transport (‘large green arrow’ in Fig.8). This establishes a cycle of mutual dependency where interruption at any point will have a knock-on effect on all other aspects of axon physiology and function (Prokop, 2021; Smith et al., 2023). Apart from explaining MT-curling in the context of this work, this cycle can explain MT-curling upon loss of other groups of genes (Hahn et al., 2021; Qu et al., 2022) and offers potential explanations for why the same classes of axonopathies can be caused by a wide range of genes involved in very different cellular functions (Prokop, 2021; Shields et al., 2024; Smith et al., 2023).

**Fig. 8.**
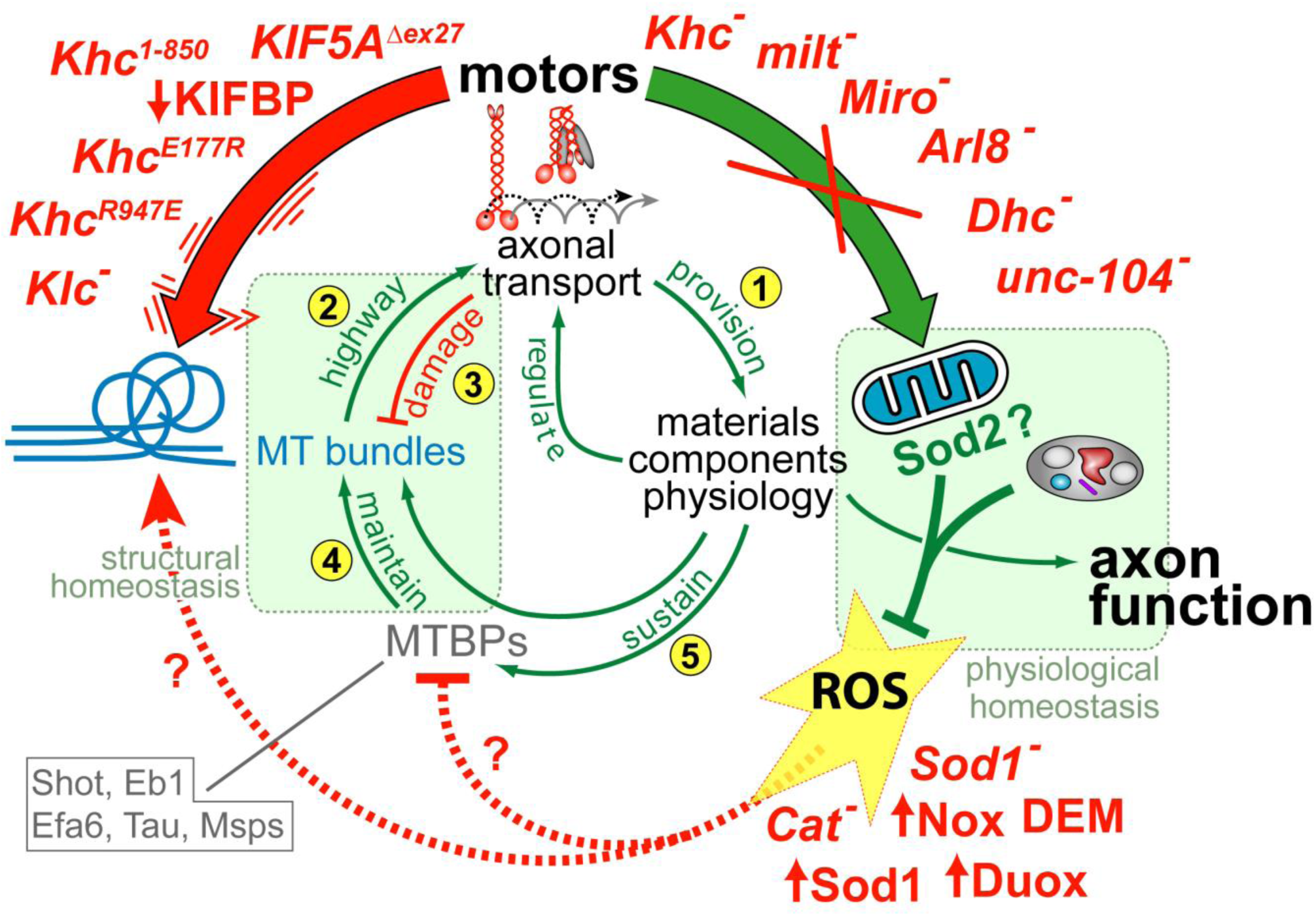
Mapping the findings of this work on the dependency cycle of local axon homeostasis. The numbered green arrows and red T-bar make up the previously published ’dependency cycle of local axon homeostasis’ (Prokop, 2021; Shields et al., 2024; Smith et al., 2023): 1) axonal transport provides materials, components and organelles required for axon function; 2) this transport requires MT bundles as the essential highways; 3) this live-sustaining transport damages MT bundles; 4) MT-binding proteins (MTBPs) support and maintain MT bundles (emboxed names in grey are proteins shown to be involved in MT bundle-maintenance; Alves-Silva et al., 2012; Hahn et al., 2021; Qu et al., 2019); 5) bundle maintenance depends on transport of components and physiology, thus closing the circle. Disturbing physiological homeostasis (light green box on the right) causes MT-curl-inducing ROS, whereas skipping the balance between motor forces and MTBP-dependent maintenance mechanisms disturbs structural homeostasis (light green box on the left) causing MT curling more directly. In this model, hyperactivation of motor proteins (vibrating red arrow accompanied by genetic manipulations used here) disturbs structural homeostasis, whereas loss of transport (large green arrow with mutant conditions listed in red) affects physiological homeostasis; upon normal transport, mitochondria and lysosome can regulate ROS levels (green T-bar). Used pharmacological and genetic manipulation of ROS that all cause MT curling (dashed red arrow and T-bar) are shown bottom right.

## Conflict of Interest

None of the authors has a conflict of interest.

## Acknowledgements

This work was made possible through support by the Biotechnology and Biological Sciences Research Council (BBSRC) to A.P. (BB/I002448/1, BB/P020151/1, BB/L000717/1, BB/M007553/1) and M.L. (BB/R016666/1), by parents to Y.-T.L., H.T. and T.M. The Fly Facility has been supported by funds from The University of Manchester and the Wellcome Trust (087742/Z/08/Z). We thank colleagues and the Bloomington *Drosophila* Stock Center (NIH P40OD018537) for providing fly stocks as mentioned in Materials and Methods.

## Methods

### Fly stocks

All human homology statements are based on information listed on flybase.org (Marygold et al., 2016; Millburn et al., 2016), all statements about genetic links to human diseases on information provided by www.omim.org (Online Mendelian Inheritance in Man®; Amberger et al., 2015). The following stocks were used in this study (reference and source provided in brackets; BL indicates Bloomington Drosophila Stock Collection): null mutant alleles (unless indicated differently) we used were

- *Arl8^e00336^* (*PBac(RB)Arl8e00336*; BDSC #17846; Rosa-Ferreira et al., 2018)
- *Arl8^Df^* (*Df(3R)D7*; BDSC #1898; Rosa-Ferreira et al., 2018)
- *Cat^n1^* (Mackay and Bewley, 1989; Matthias Landgraf)
- *Dhc64C^4-19^* (Gepner et al., 1996; BDSC #5274)
- *Khc^8^* (Saxton et al., 1991; BDSC #1607)
- *Khc^27^* (Saxton et al., 1991; Isabel Palacios)
- *Khc^1ts^* (Saxton et al., 1991; BDSC #31994; a temperature-sensitive allele which is homozygous viable at 18⁰C but usually kept over balancer)
- *Khc^mutA^* (Winding et al., 2016; Vladimir Gelfand; confirmed by lethality of hetero-allelic *Khc^mutA/8^*animals)
- *Khc^E177K^* and *Khc^E177K,R947E^* (Kelliher et al., 2018; Jill Wildonger)
- *Df(Khc)* (*Df(2R)BSC309*; Cook et al., 2012; BDSC #23692)
- *Klc^8ex94^* (Gindhart et al., 1998; BDSC #31997)
- *Df(Klc)* (Bloomington #7596)
- *Klp64D^k1^* (Ray et al., 1999; hypomorphic allele; BDSC #5578)
- *Klp64D^n123^* (Perez and Steller, 1996; BDSC #5674)
- *Klp98A^Δ47^* (Derivery et al., 2015; Marcos Gonzalez-Gaitan)
- *milt^92^* (Cox and Spradling, 2006; Stowers et al., 2002; Tom Schwarz)
- *Df(milt)* (*Df(2L)ED440, P{w[+mW.Scer\FRT.hs3]=3’.RS5+3.3’}ED440*; Ryder et al., 2004; Kyoto #150498)
- *Miro^Sd32^* (Guo et al., 2005)
- *Miro^B682^* (Guo et al., 2005; BDSC #52003)
- *Df(Miro)* (*Df(3R)Exel6197*; Parks et al., 2004; BDSC #7676)
- *Pat1^grive^* and *Pat1^robin^* (Loiseau et al., 2010; Isabel Palacios)
- *Pex3^2^* (Faust et al., 2014; BDSC #64251)
- *Sod1^n1^* (Phillips et al., 1989; BDSC #24492)
- *Sod1^n64^* (Phillips et al., 1995; BDSC #7451)
- *syd^z4^* (Bowman et al., 2000; BDSC #32016)
- *Df(3L)syd^A2^* (Bowman et al., 2000; deleting C-terminus; BDSC #32017)
- *unc-104^170^* (Pack-Chung et al., 2007; Tom Schwarz)

Gal4 driver lines used were the

- *elav-Gal4* (BDSC #8765)
- *tubP-Gal4* (Lee and Luo, 1999; Liqun Luo)

UAS lines

- *UAS-Duox* (Ha et al., 2005; Matthias Landgraf)
- *UAS-Gapdh-IR* (*Gapdh1^GD7467^*; VDRC #31631)
- *UAS-Khc^FL^::GFP* (3^rd^, unpublished; Isabel Palacios)
- *UAS-Khc1-850-GFP* (Loiseau et al., 2010; Isabel Palacios)
- *UAS-Khc-RNAi* (Lu et al., 2013; Vagnoni et al., 2016; BDSC #35770)
- *UAS-hKIF5A^wt^-GFP* (Pant et al., 2022; Devesh Pant)
- *UAS-hKIF5A^Δexon27^-GFP* (Pant et al., 2022; Devesh Pant)
- UAS-KIFBP-IR (CG14043; BDSC #51858; Perkins et al., 2015)
- *UAS-Nox* (Liew et al., 2021)
- *UAS-Sod1* (J. Hu and J.P. Phillips, unpublished)
- *UAS-YFP-PTS1* (BDSC #64248; Faust et al., 2014)

### Cloning of *UAS-Nox-YPet*

*10xUAS-IVS-Nox::YPet* was generated by using the *pJFRC12-10XUAS-IVS-myr-GFP* vector (Addgene 26222; Pfeiffer et al., 2010) as a backbone which was modified by substituting GFP with YPet (Nguyen and Daugherty, 2005) plus an N-terminal flexible linker, amplified from dFlex_YPet_phase0 (Gärtig et al., 2019) using primer ML1 and ML2, and inserted by the Klenow Assembly Method (https://tinyurl.com/4r99uv8m) into the XbaI/BamHI sites producing Vector 1: *pJFRC12-10xUAS-IVS-myr-linker-YPet*. Nox cDNA was amplified from a DGRC (*Drosophila* Genomics Resource Center) cDNA library clone using primers ML5 and ML6, located in a *pOTB7* vector backbone, and inserted into BamHI/XhoI sites of Vector 1. Constructs were sent to FlyORF for transgenesis, and targeted via PhiC31-mediated site-specific insertion to the *PBac{y^+^-attP-3B}VK00040* landing site (Bloomington line #9755) on the third chromosome (3R, 87B10).

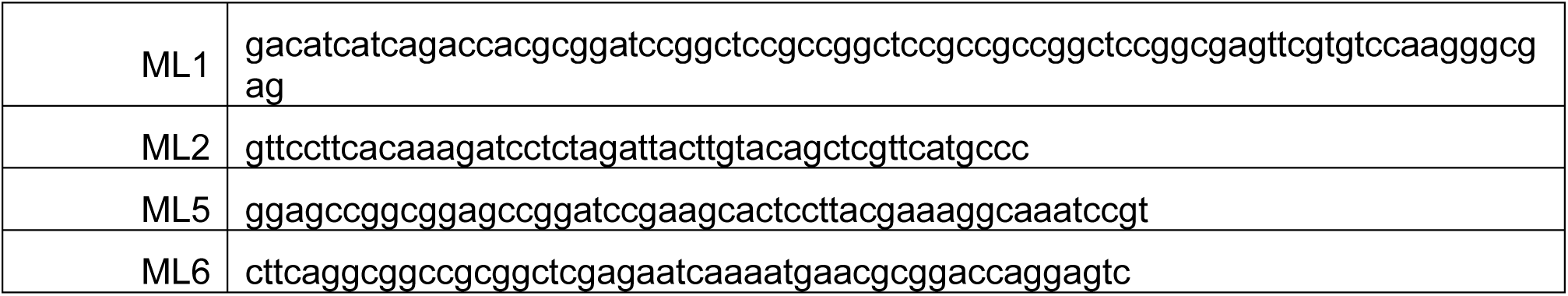

### *Drosophila* primary cell culture

*Drosophila* primary neuron cultures were performed as published previously (Prokop et al., 2012; Qu et al., 2017). In brief, stage 11 embryos were treated for 1 min with bleach to remove the chorion, sterilized for ∼30 s in 70% ethanol, washed in sterile Schneider’s/FCS, and eventually homogenized with micro-pestles in 1.5 centrifuge tubes containing 21 embryos per 100μl dispersion medium and left to incubate for 5 min at 37°C. Cells were washed with Schneider’s medium (Gibco), spun down for 4 mins at 650g, supernatant was removed and cells re-suspended in 90µl of Schneider’s medium containing 20% fetal calf serum (Gibco). 30μl drops were placed on cover slips. Cells were allowed to adhere for ∼2hrs either directly on glass or on cover slips coated with a 5 µg/ml solution of concanavalin A, and then grown as a hanging drop culture for hours or days at 26°C as indicated in each experiment.

To abolish maternal rescue of mutants, i.e. masking of the mutant phenotype caused by deposition of normal gene product from the healthy gene copy of the heterozygous mothers in the oocyte (Prokop, 2013), we used a pre-culture strategy (Prokop et al., 2012; Sánchez-Soriano et al., 2010) where cells were kept for 5 days in a tube before they were plated on a coverslip.

Cells were treated with 100 µM Trolox (97% purity, Thermo Scientific or Sigma; stepwise diluted from a 100 mM stock solution in ethanol) in 100% ethanol. For controls (vehicle treatment), equivalent concentrations of vehicle (100% ethanol) were diluted in cell culture medium. In later experiments, cells were treated with 100μM Trolox or 100μM diethyl maleate (DEM) dissolved directly into neuronal culture medium; where necessary, the solution was incubated at 26 °C and periodically agitated to ensure it is fully dissolved. The solution was then filter-sterilised. The treated cells were grown in this medium for the entire culture period whereas control cells were grown in Trolox/DEM-free medium. All reagents were purchased from Sigma-Aldrich, unless otherwise stated.

For visualisation of mitochondria, cell cultures were incubated with 400nM MitoTracker Red CMXRos (Invitrogen; Klionsky et al., 2012) for 30min at room temperature (RT) before fixation; stock solutions were prepared in DMSO and diluted in cell culture medium to the final concentration. Following incubation, cultures were then fixed and stained following the procedures below.

### Immunohistochemistry

Primary fly neurons were fixed in 4% paraformaldehyde (PFA) in 0.05M phosphate buffer (PB; pH 7–7.2) for 30min at room temperature (RT). Antibody staining and washes were performed with PBT. Staining reagents: anti-tubulin (clone DM1A, mouse, 1:1000, Sigma; alternatively, clone YL1/2, rat, 1:500, Millipore Bioscience Research Reagents); anti-Syt (1:1000; rabbit; Sean Sweeney); anti-GFP (1:500, rabbit, ab290, Abcam); Cy3-conjugated anti-HRP (goat, 1:100, Jackson ImmunoResearch); FITC-, Cy3- or Cy5-conjugated secondary antibodies (1:200; donkey, purified, Jackson Immuno Research); F-actin was stained with Phalloidin conjugated with TRITC/Alexa647, FITC or Atto647N (1:200; Invitrogen and Sigma). Specimens were embedded in ProLong Gold Antifade mounting medium.

### Mouse cell culture

Mice (C57BL/6) were housed, bred, and treated in compliance with the Animal (Scientific Procedures) Act 1986 at the Biological Support Unit, University of Nottingham. Primary cortical neuron cultures were prepared from time-mated embryos at embryonic day 16.5 (E16.5). The brains were removed, placed in Hanks’ Balanced Salt Solution (HBSS; Sigma), and the cortices isolated under a dissection microscope, with meninges removed using forceps. The tissue was transferred to a 35mm Nunc™ dish with 2 ml of HBSS and mechanically dissociated. Trypsin (1 mg/ml; Sigma-Aldrich) and DNAse I (50 µg/ml; Sigma-Aldrich) were added, followed by incubation at 37°C (5% CO_2_) for 30 minutes. Soybean trypsin inhibitor (0.05%; Sigma-Aldrich) was then added. The tissue was mechanically dissociated further, and cells were re-suspended in Neurobasal^®^ medium (Gibco) with 1% GlutaMAX™ (Gibco) and 2% B-27™ supplement (Gibco). Cells were plated at a 1:144 dilution (from a stock solution of 10-12 x 10^6^ cells/ml) on 12-well plates (Corning) with 19 mm glass coverslips (Fisher Scientific) coated with 50 µg/ml poly-L-ornithine (Sigma-Aldrich), incubated overnight, and washed twice with dH2O before use. 100μM or 200μM of diethyl maleate (DEM; Sigma) were dissolved directly in culture medium, which was then filter-sterilised and added to the cultured cells on day 3.

4% PFA in PBS (Thermo Scientific™) was used to fix the neurons on day 4, 24 hours after DEM application. The cell culture medium was discarded, and coverslips were rinsed with PBS before being incubated in PFA for 30 minutes at room temperature. The PFA was discarded, and coverslips were washed with PBS (3 x 5 minutes) and Buffer 1 (10mM glycine in PBS; 3 x 5 minutes). They were then incubated in Buffer 2 (10mM glycine and 0.2% Triton™ X-100 in PBS) for 30 minutes at room temperature, followed by washing with Buffer 3 (0.1% Triton™ X-100 in PBS; 3 x 5 minutes). Coverslips were incubated in PBS with 3% BSA (Sigma-Adrich) for 30 minutes, then placed onto 25 µl droplets of primary antibody (anti-acetylated tubulin, 1:300 in BSA 3%, T7451, Sigma-Aldrich) in a humidified chamber at 4°C overnight. The next day, coverslips were washed with Buffer 3 (3 x 5 minutes) and placed onto droplets containing the secondary antibody (488-coupled goat anti-mouse, 1:300 in BSA 3%, A11001, Invitrogen) and DAPI (1:300 in BSA 3%; Sigma-Aldrich) for 2 hours at room temperature. Coverslips were then washed with Buffer 3 (3 x 5 minutes) and stored in PBS until mounting. They were mounted onto Epredia™ SuperFrost™ slides (Fisher Scientific) using 12 µl VectaShield® HardSet™ mounting media (Vector Laboratories) per coverslip and stored at 4°C until imaging.

### Microscopy and data analysis

Standard documentation was performed with AxioCam monochrome digital cameras (Carl Zeiss Ltd.) mounted on BX50WI or BX51 Olympus compound fluorescent microscopes. To determine the degree of MT disorganisation in axons we used the "MT disorganisation index" (MDI) (Qu et al., 2017): the area of disorganisation was measured using the freehand selection tool in Fiji/ImageJ; this value was then divided by axon length (see above) multiplied by 0.5 μm (typical axon diameter, thus approximating the expected area of the axon if it were not disorganised). To quantify the number of synaptic densities in mature neurons in culture, we used ImageJ, first thresholding to select synaptic densities from axons of single isolated cells, followed by particle analysis. For statistical analyses, Kruskal–Wallis one-way ANOVA with *post hoc* Dunn’s test or Mann–Whitney Rank Sum Tests were used to compare groups. The data used for our analyses will be made available on request from the authors. Experiments were repeated at least once, data pooled. MDI data were usually not normally distributed but nevertheless plotted as mean ± SEM to avoid misleading median values of zero. Most experiments had only two groups and were assessed using Mann–Whitney Rank Sum tests, for experiments with more then two groups we applied Kruskal–Wallis one-way ANOVA with *post hoc* Dunn’s test. Means of single slides were used to generate super-plots (Lord et al., 2020) and assessed using standard t-tests. The data used for our analyses will be made available on request.

### Expression analysis of KIF5AΔex27::GFP

HEK293T cells were maintained in Corning® DMEM (Dulbecco’s Modified Eagle’s Medium, Cat# 45000-304) supplemented with 10% fetal bovine serum (Cat# 35-011-CV) and 1× penicillin-streptomycin-glutamine (Cat# 10378016). Cells were cultured at 37°C in a humidified incubator with 5% CO₂. For transient transfection, cells were transfected using polyethylenimine (1 mg/mL) with the human *KIF5A^Δex27^-GFP* plasmid and incubated for 48 hrs. At 24 hrs post-transfection, cells were treated with 1.5 mM Trolox (Cat# 218940050) for the remaining 24 hours.

Whole-cell extracts were prepared using 8 M urea buffer (10 mM Tris, pH 8.0; 8 M urea) supplemented with the Halt™ protease and phosphatase inhibitor cocktail (Cat# PI78441). Protein concentration was determined using the BCA Protein Assay Reagent (Cat# PI23227). Urea protein lysates were resolved on a 4–20% precast polyacrylamide gel (Bio-Rad, USA) and transferred to nitrocellulose membranes. Membranes were incubated overnight at 4°C with primary antibodies: mouse anti-GFP (1:1,000; Cat# 632381) and rabbit anti-ACTIN (1:1,000; Cat# GTX637675). This was followed by incubation with HRP-conjugated secondary antibodies (Cat# AS003) or IRDye secondary antibodies (Cat# 926-32211) for 1 hour at room temperature. Signal detection was performed using SuperSignal™ West Pico Chemiluminescent Substrate (Cat# PI34578). Molecular weights were assessed by comparison to a protein ladder (Cat# 26619), and band intensities were quantified using ImageJ, normalized to actin.

## Ethical statement

An ethical statement is not required.

## Figures

**Fig. S1.**
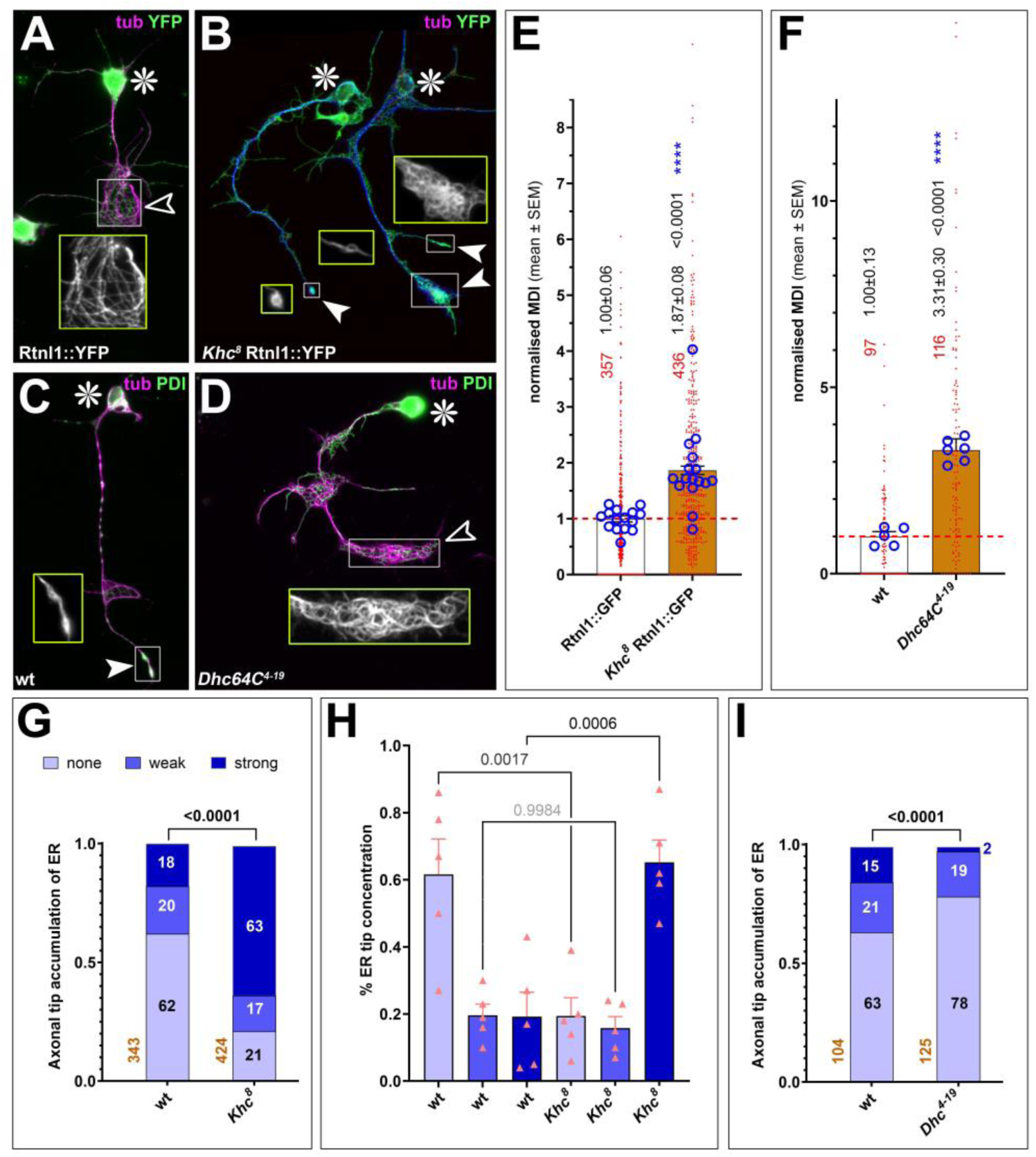
Upon loss of Khc, the endoplasmatic reticulum tends to accumulate at axon tips, independent of MT curling. **A-D**) Primary neurons at 5 DIV either being wild-type (A,C) or homozygous mutant for the *Khc^8^* (B) or *Dhc64C^4-19^* (D) mutant alleles. MTs are labelled in magenta (tub) and the endoplasmic reticulum (ER) in green: either via fluorescence of the genomically tagged *Rtln1-YFP* allele (del Castillo et al., 2019; O’Sullivan et al., 2012) or using anti-PDI staining; white boxes are shown as 2-fold magnified, yellow-framed insets showing only the MTs; cell bodies are indicated by asterisks, axon tips with/without strong ER accumulation by white/open arrowheads; as can be seen, there is no correlation between MT curling and ER accumulation. **E,F**) Control measurements indicate that the MT-curling phenotypes are reliably present in the neurons analysed for ER tip accumulation (for further explanation of graphs see legend of Fig.1); note that analyses of *Khc^8^* Rtnl1::YFP mutant neurons were repeated in 2016, 2017 and 2022 with reproducible results (see also H). **G,I**) Quantification of no/weak/strong ER accumulation at axonal tips (indicated by light/medium/dark blue) in Rtnl1::YFP or wt controls and in *Khc^8^* Rtnl1::YFP or *Dhc64C^4-19^* mutant neurons; the P value of Х^2^ tests is shown above bars, the number of analysed neurons indicated in red. **H**) The pink triangles differently display the data from G representing the respective percentages of 5 independent experiment between 2016 and 2022 compared by t-tests, indicating a consistent increase in tip localisation upon loss of Khc. This increase is surprising contradicting reports in mouse neurons (Farías et al., 2019). We reasoned this to be caused by loss of Khc-dependent Dynein transport to axon tips (Moughamian et al., 2013; Twelvetrees et al., 2016), but loss of Dhc unexpectedly causes the opposite phenotype.

**Fig. S2.**
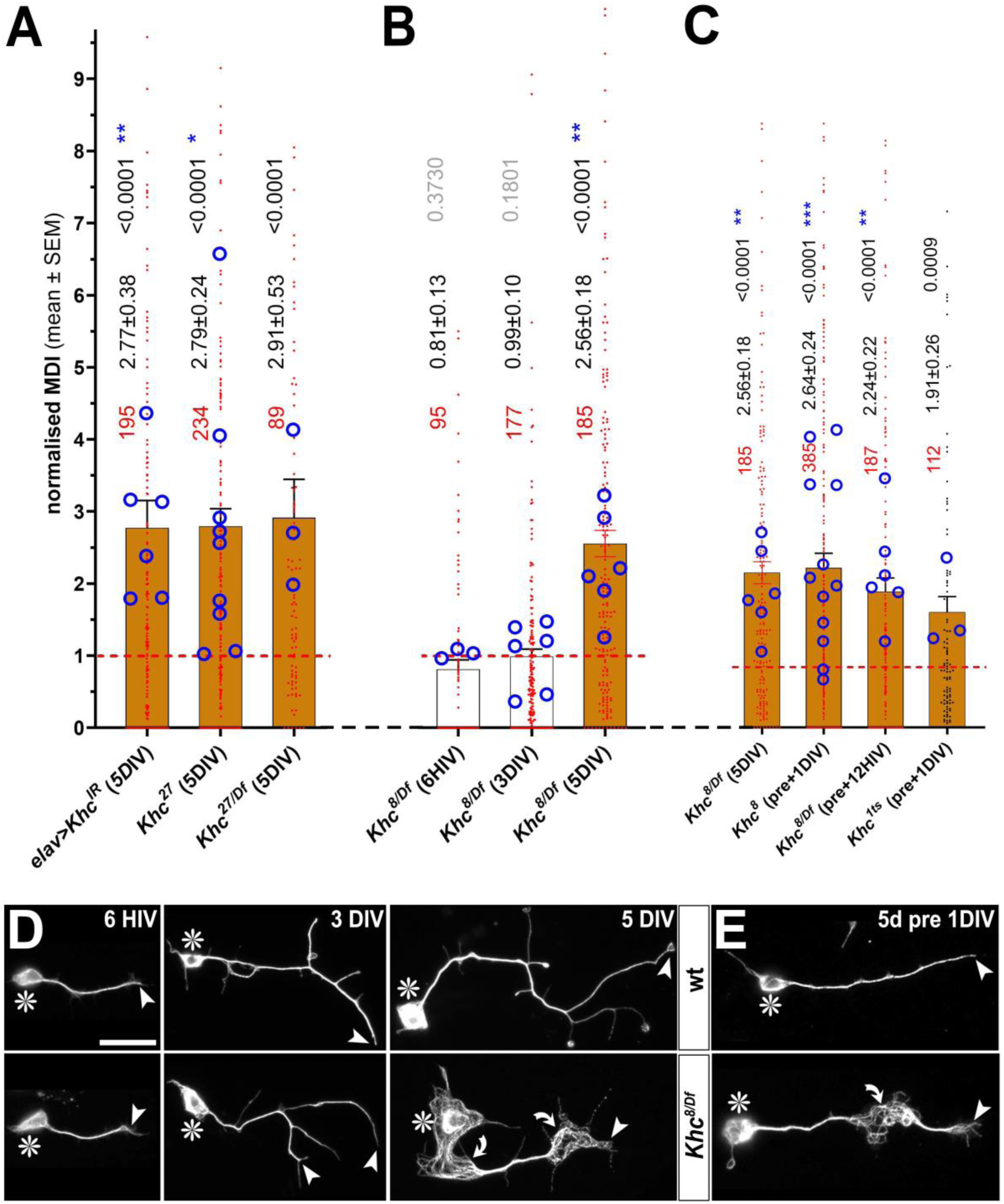
Validation of Khc’s MT phenotype and demonstration of maternal contribution. **A-C**) Quantification of MT-curling phenotypes measured as MT disorganisation index (MDI) and normalised to internal wild-type of *elav-Gal4/+* controls (red stippled line). A) shows data for *Khc^27^*in homozygosis (*27*) or over deficiency (*27/Df*), for Khc knock-down (*elav>Khc-IR*). B) shows data for *Khc^8^* over deficiency (*8/Df*) at different culture times (HIV, hours *in vitro*; DIV, days *in vitro*); C) shows data for *Klc^8/Df^* and *Klc^1ts^*at 12HIV or 1DIV following 5d pre-culture; note that *Klc^1ts^* is a temperature-sensitive allele (see methods) and was pre-cultured at 26⁰C and cultured at 29⁰C. For further graph descriptions refer to the legend of Fig.1. **D,E**) Examples of neurons at different times in culture (D; relating to data in B) and after pre-culture (E; relating to C); asterisks indicate cell bodies, arrow heads axon tips, curved arrows areas of MT curling; scale bar in D represents 20µm in D and E.

**Fig. S3.**
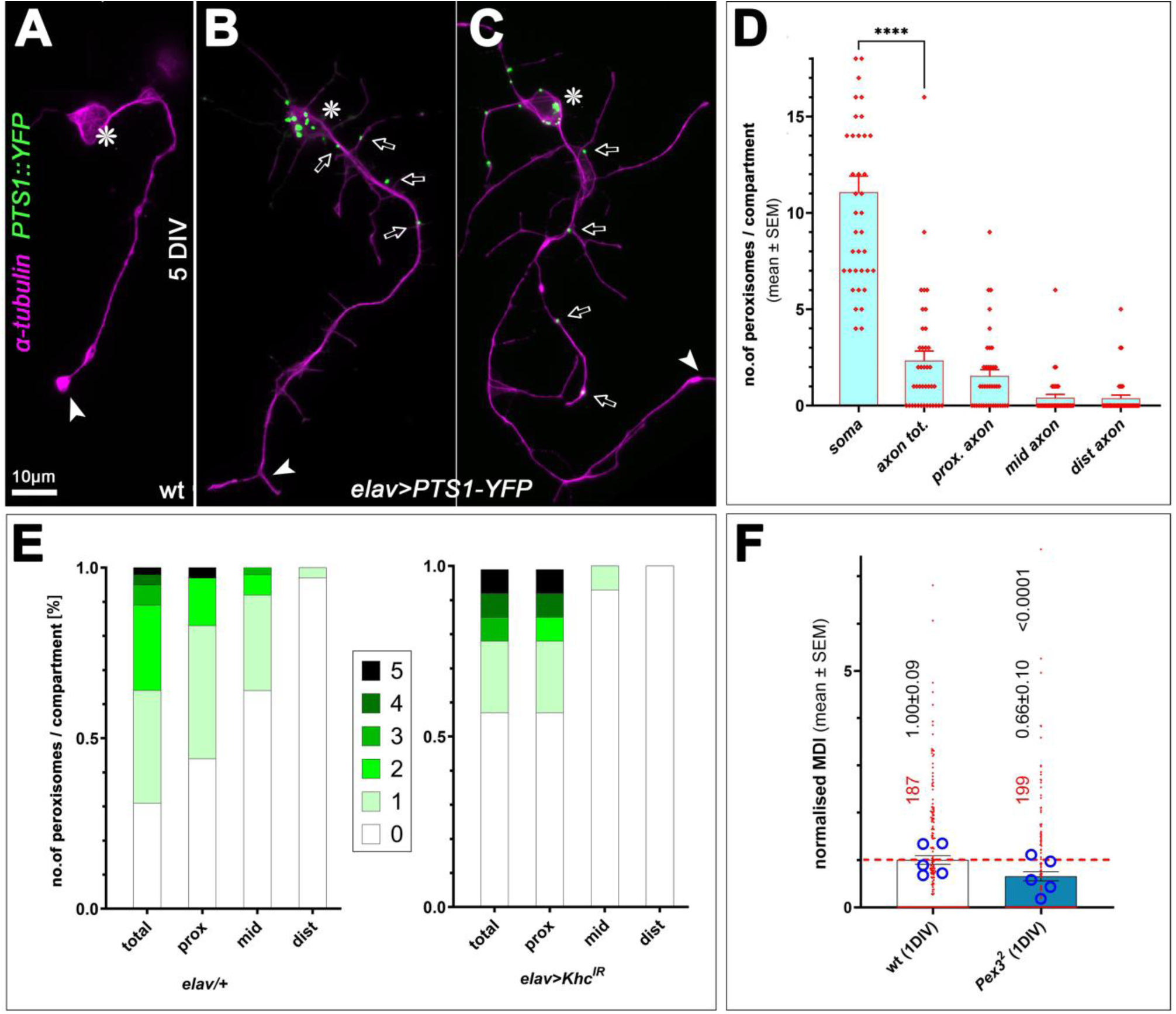
Analyses of peroxisomes in *Drosophila* primary neurons. **A-C)** Images of *Drosophila* primary neurons at 5DIV without (A) or with (B,C) expression of PTS1::YFP, stained for tubulin (magenta); asterisks indicate cell bodies, arrowheads tips of axons; arrows point at labelled peroxisomes; scale bar in A represents 10µm in all images. **D)** Quantification of peroxisome numbers classified by cell compartments: soma, total axon, proximal axon, mid axon and distal axon. **E)** Distribution of peroxisomes in axons in either controls (*elav/+*) or upon Khc knock-down (*elav>Khc^IR^*). **F)** Quantification of MT-curling in Pex3-deficient neurons; for explanations of the graph structure, refer to Fig.1.

**Fig. S4.**
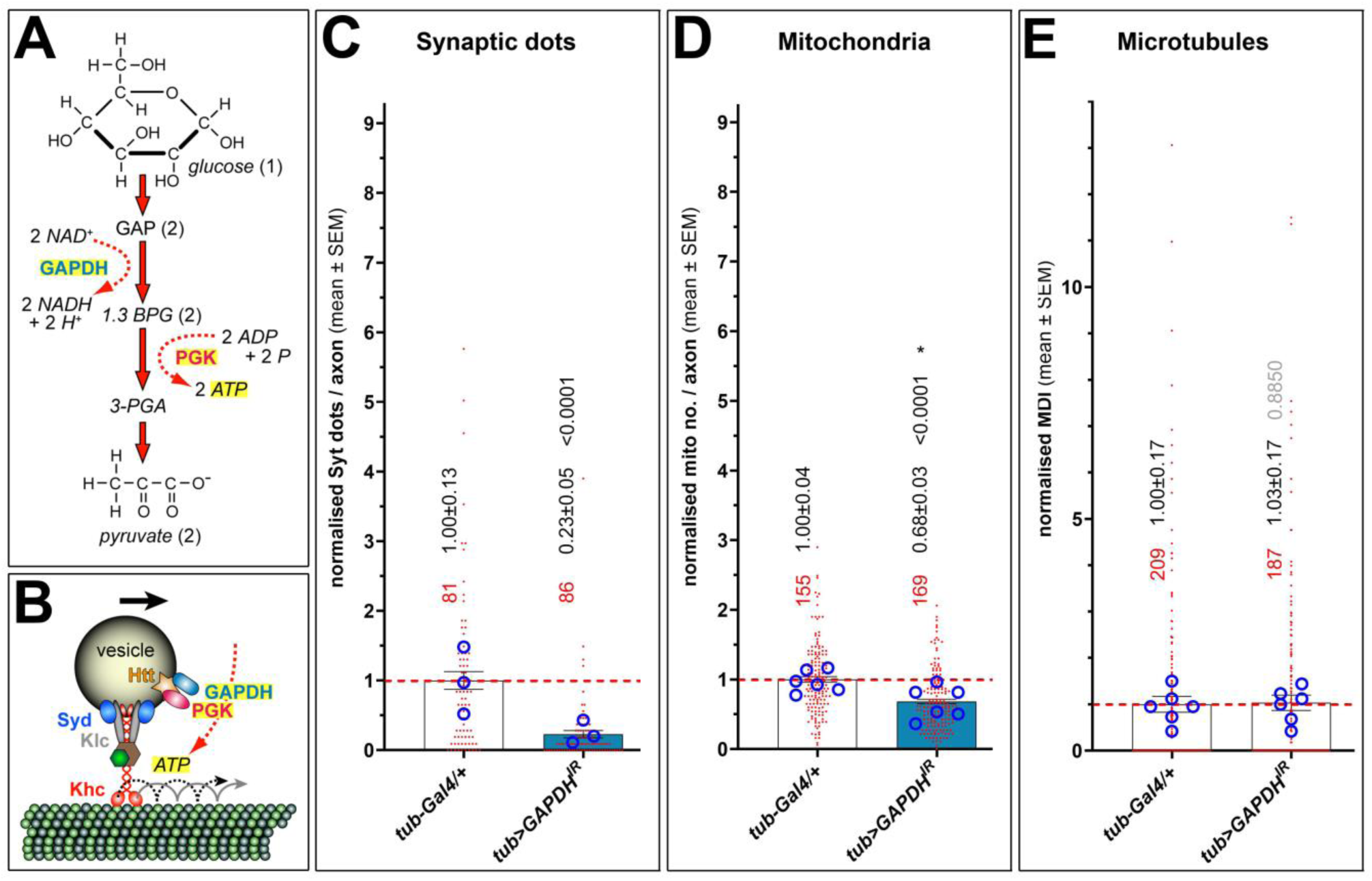
Phenotypes upon *Gapdh1* knock-down in primary neurons at 5 DIV. **A**) Illustration of the NADH- and ATP-generating steps of glycolysis; names of proteins are shown in bold, other molecules in italics: GAPDH (glyceraldehyde-3-phosphate dehydrogenase), PGK (phosphoglycerate kinase), GAP (glyceraldehyde-3-phosphate), 1,3 BPG (1,3-biphosphoglycerate), 3-PGA (3-phosphoglycerate). **B**) GAPDH and PGK are present on transported vesicles together with other factors relevant for glycolysis (Hinckelmann et al., 2016; Zala et al., 2013) providing ATP to drive kinesin-mediated processive transport (red and stippled black lines). **C-D**) In the absence of Gapdh1, the transport of synaptic vesicles but not mitochondria is impaired (assessed via anti-Syt and mitoTracker staining; see Fig.2). **E**) Absence of GAPDH does not cause MT curling. For explanations of graph organisation, refer to the legend of Fig.1.

**Fig. S5.**
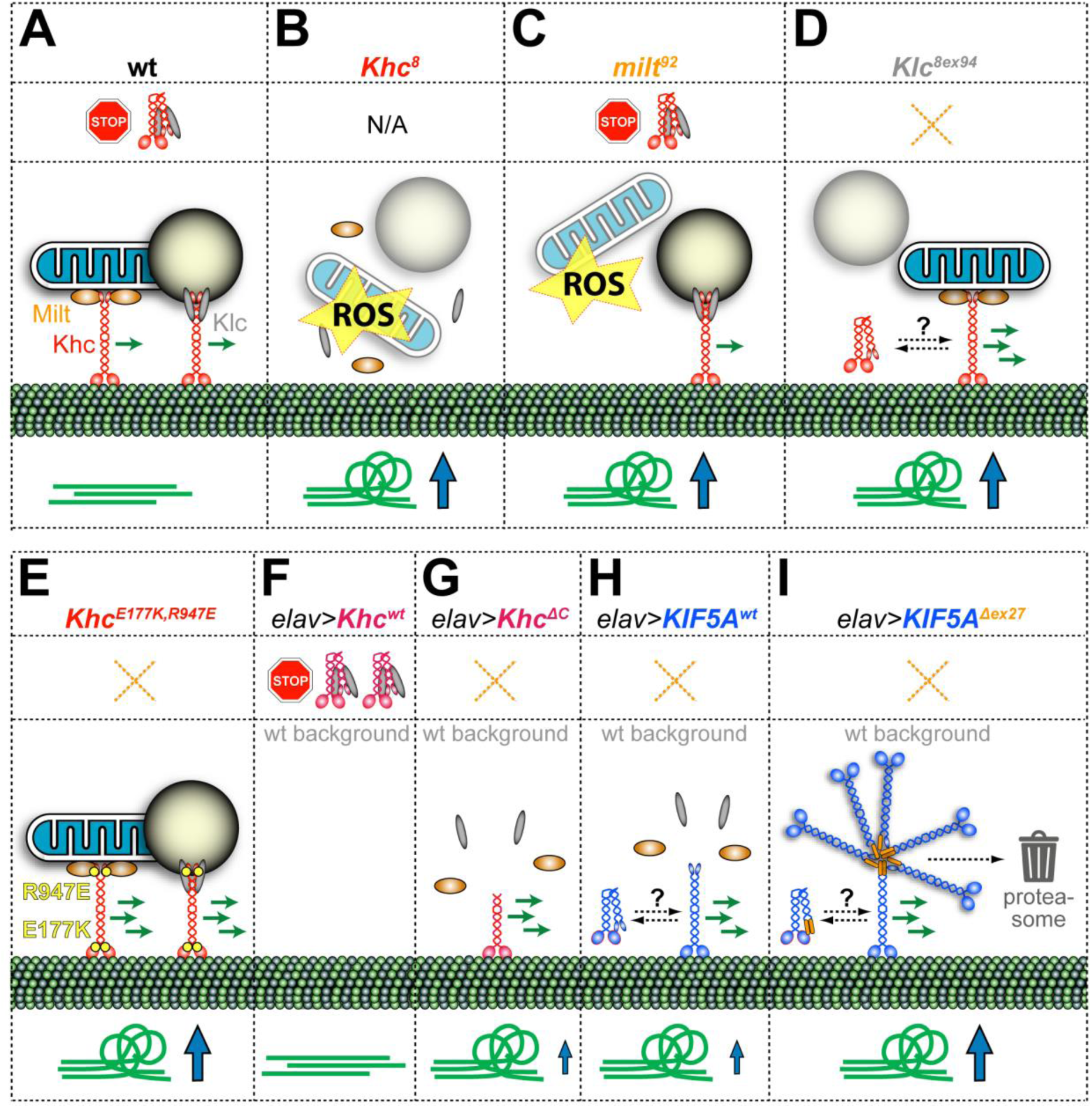
Graphical summary of key findings in this paper. A-I each represent one genotype: in each case, the top section lists the genotype, the second section indicates whether this condition permits auto-inhibition (N/A, not applicable), the third section lists the impact on transport (the number of green arrows reflects the amount of transport, organelles with low transparency indicate affected transport) or harmful ROS production, the bottom section indicates whether MT-curling is increased (size of blue arrow reflects the degree).

**Fig. S6.**
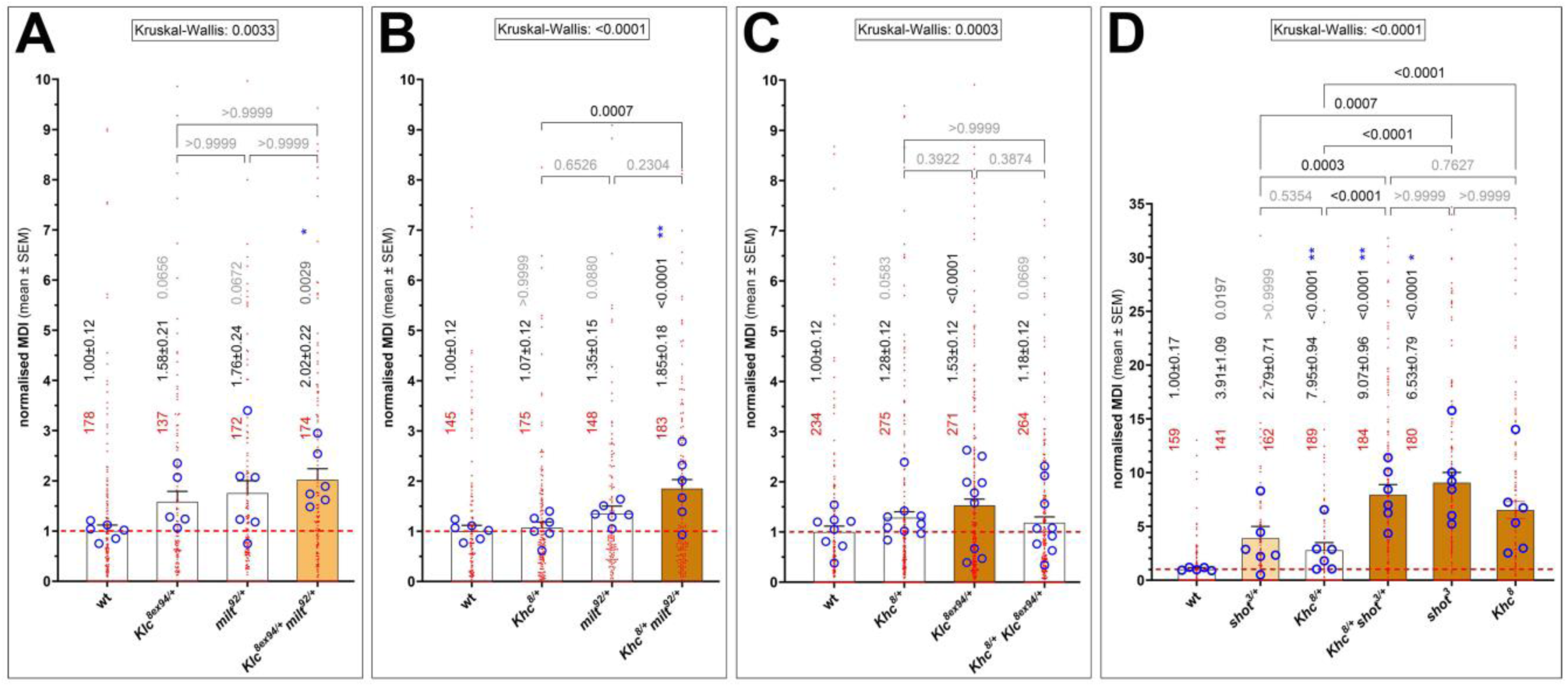
Genetic interaction studies involving Khc, Milt and Klc. Each graph represents a separate genetic interaction experiment showing wild-type controls, the two single-heterozygous conditions followed by the trans-heterozygous condition; in D also the homozygous mutant conditions for *shot^3^*and *Khc^8^* are shown. For further explanations of graph organisation, refer to the legend of Fig.1.

**Fig. S7.**
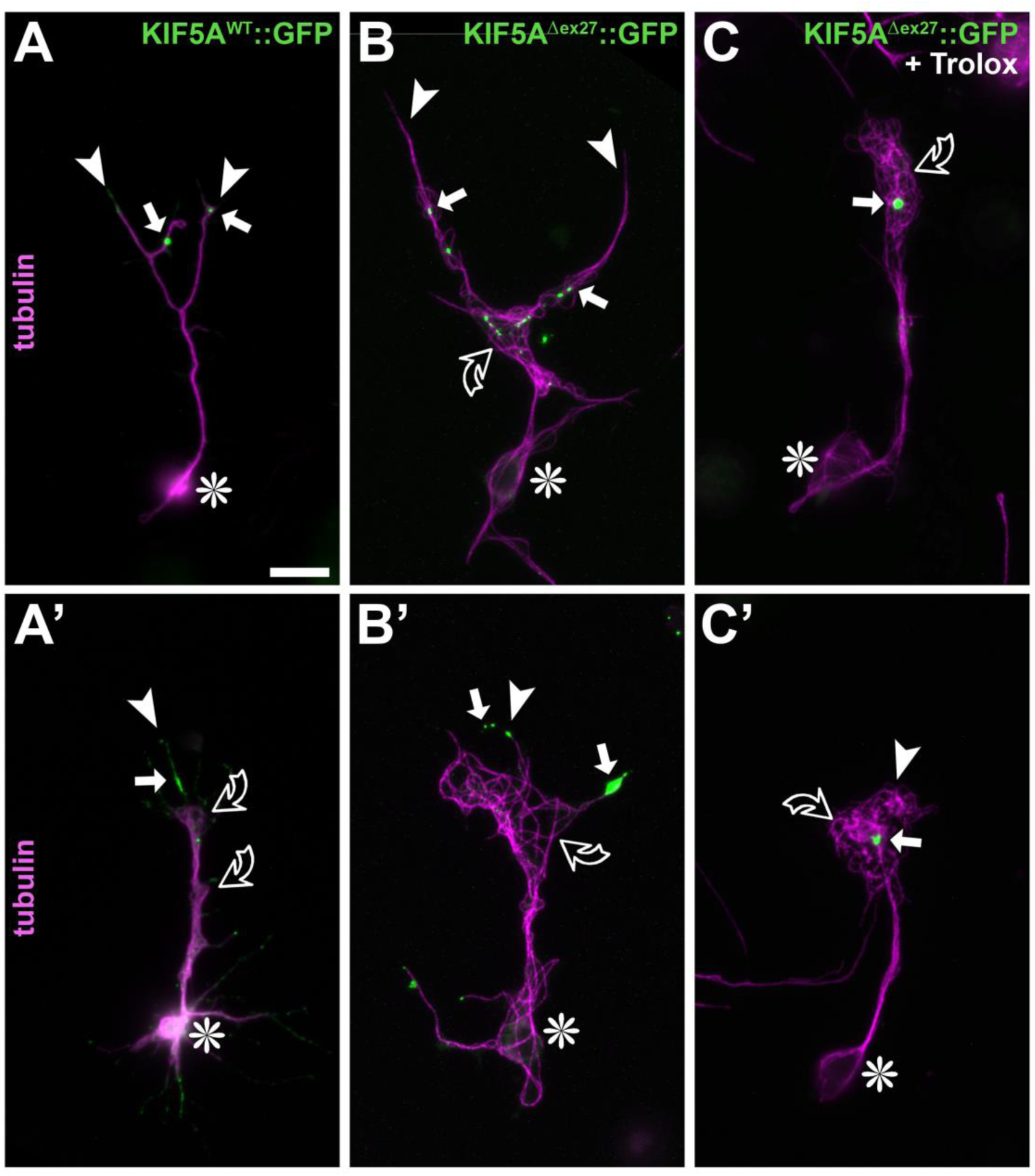
Examples of neurons expressing normal and mutant versions of human KIF5A. *Drosophila* primary neurons at 3DIV expressing *elav-Gal4*-driven KIF5A^wt^::GFP or KIF5A^Δex27^::GFP (as indicated top right) and stained for tubulin, treated with Trolox in C,C’; asterisks indicate cell bodies, arrow heads axon tips, open curved arrows areas of MT curling, white arrow distal GFP aggregates; scale bar in A represents 20µm in all images.

**Fig. S8.**
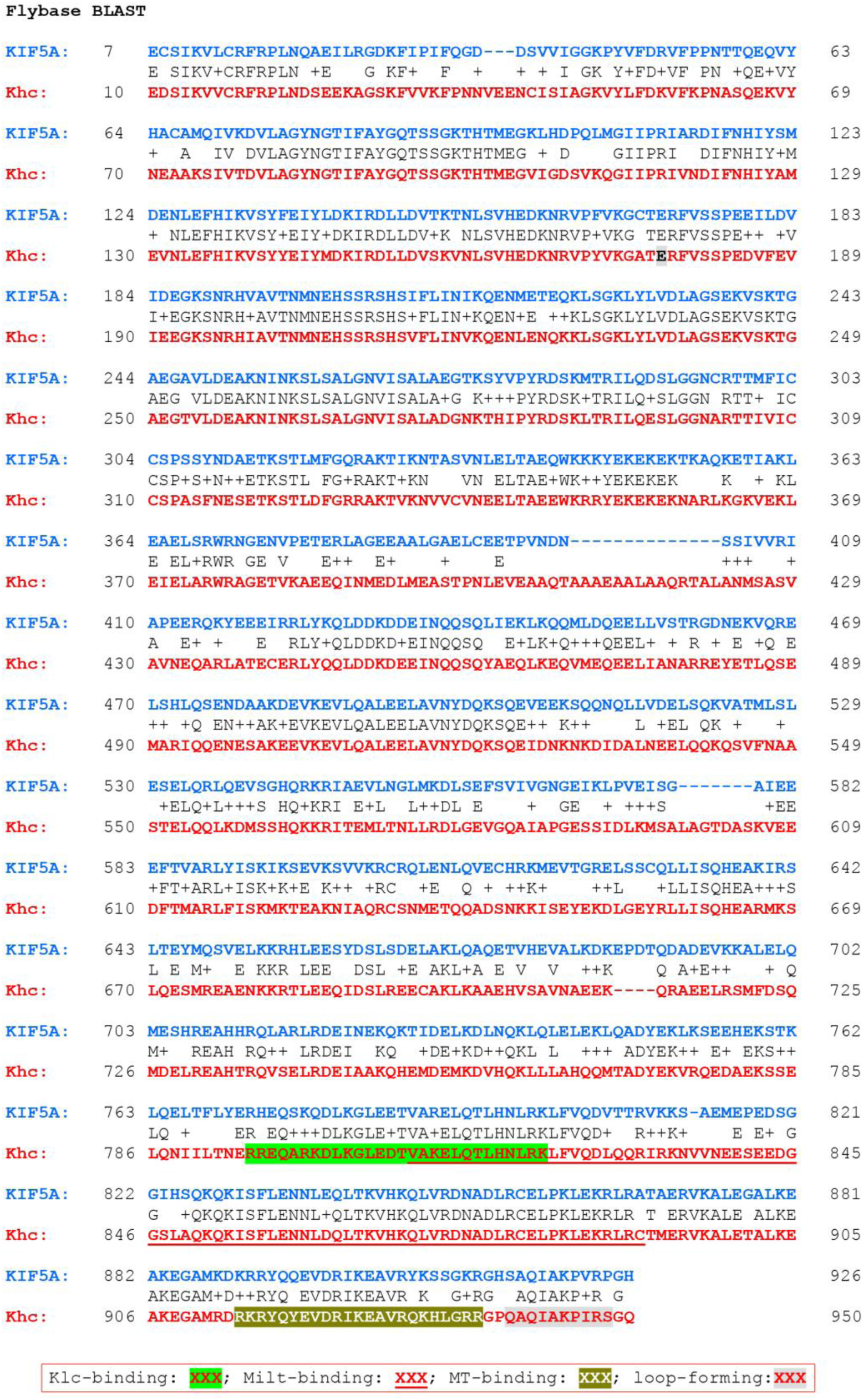
Sequence alignment of KIF5A and Khc. Alignment of KIF5A-202 (ENST00000455537.7, 5779bp,1032aa) and Khc-PA (FBtr0087184, 3718bp, 975aa) according to the BLAST function of flybase.org. C-terminal functional regions of Khc indicated in Fig.3A are highlighted as indicated in the box at the bottom. Degree of identity: 562 / 945 (59.5%); degree of similarity: 718 / 945 (76%); gaps = 29 / 945 (3.1%).

**Fig. S9.**
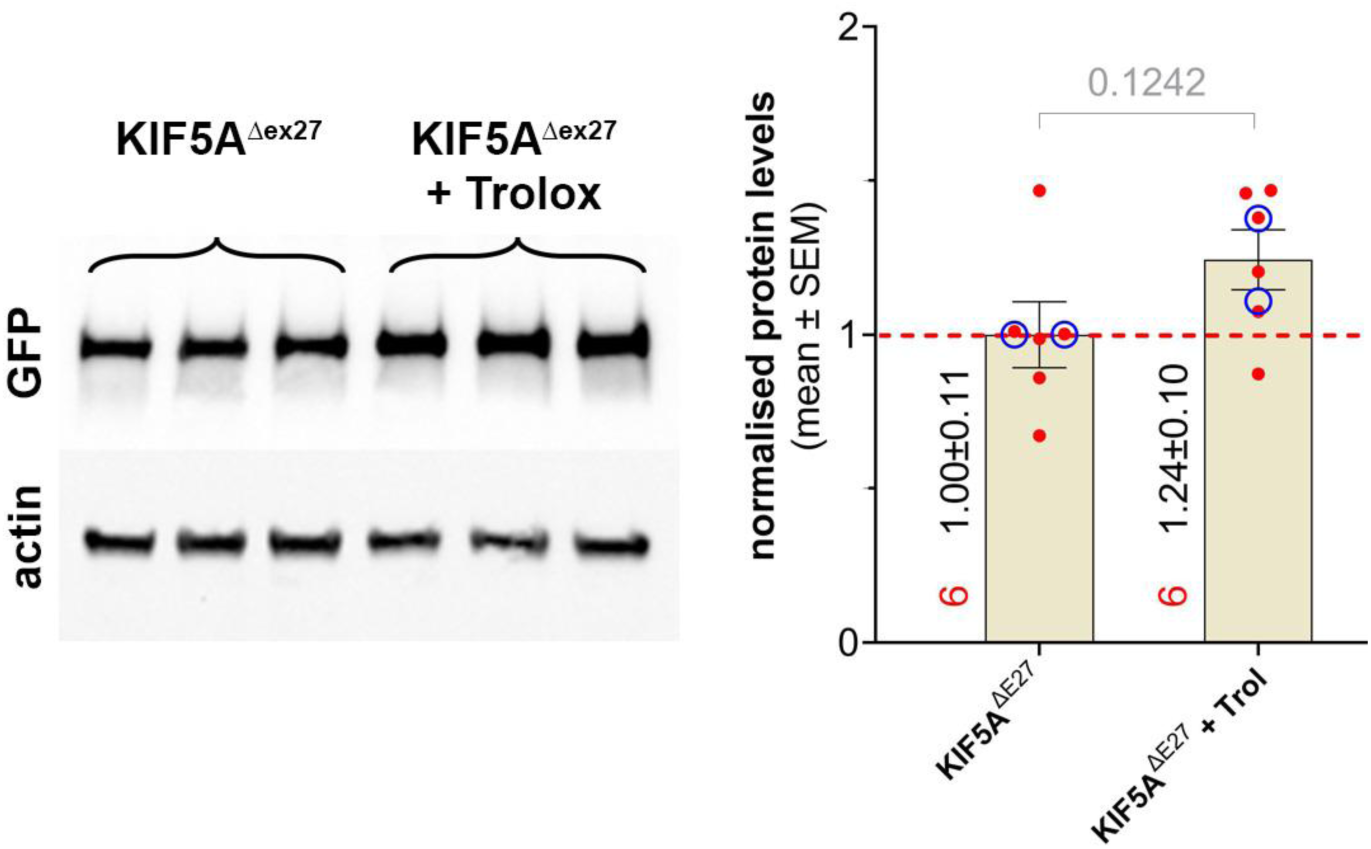
KIF5AΔex27 expression in HEK293T cells shows a trend to increased protein levels upon Trolox treatment. **Left:** HEK293T cells were transfected with *KIF5A^Δex27^-GFP* for 48 hrs and treated with 1.5 mM Trolox during the last 24 hrs; protein was prepared in 8M urea buffer, resolved on a polyacrylamide gel, blotted and stained for GFP and actin. **Right:** Graph representing the measured protein levels from 2 replicates (blue rings) with 3 repeats, respectively (red dots).

## Notes

### Competing Interest Statement

The authors have declared no competing interest.

### Summary of Updates

Additional authors additional data in Figs. 2, 6, 7, S2, new figure S9 improvement to the text throughout and to some images New data support the wider application of our results to other transport motor proteins beyond kinesin-1

